# Preferential sampling for presence/absence data and for fusion of presence/absence data with presence-only data

**DOI:** 10.1101/409680

**Authors:** Alan. E. Gelfand, Shinichiro Shirota

## Abstract

Presence/absence data and presence-only data are the two customary sources for learning about species distributions over a region. We present an ambitious agenda with regard to such data. We illuminate the fundamental modeling differences between the two types of data. Most simply, locations are considered as fixed under presence/absence data; locations are random under presence-only data. The definition of “probability of presence” is incompatible between the two. So, we take issue with modeling strategies in the literature which ignore this incompatibility, which assume that presence/absence modeling can be induced from presence-only specifications and therefore, that fusion of presence-only and presence/absence data sources is routine.

We argue that presence/absence data should be modeled at point level. That is, we need to specify a surface which provides the probability of presence at any location in the region. A realization from this surface is a binary map yielding the results of Bernoulli trials across all locations; this surface is only partially observed. Presence-only data should be modeled as a (partially observed) point pattern, arising from a random number of individuals at random locations, driven by specification of an intensity function. There is no notion of Bernoulli trials; events are associated with areas.

We further argue that, with just presence/absence data, preferential sampling, using a shared process perspective, can improve our estimated presence/absence surface and prediction of presence. We also argue that preferential sampling can enable a probabilistically coherent fusion of the two data types.

We illustrate with two real datasets, one presence/absence, one presence-only for invasive species presence in New England in the United States. We demonstrate that potential bias in sampling locations can affect inference with regard to presence/absence and show that inference can be improved with preferential sampling ideas. We also provide a probabilistically coherent fusion of the two datasets to again improve inference with regard to presence/absence.

The importance of our work is to provide more careful modeling when studying species distributions. Ignoring incompatibility between data types and offering incoherent modeling specifications implies invalid inference; the community should benefit from this recognition.

## 1 Introduction

Learning about species distributions is, arguably, a preoccupation in the ecology community. The literature discusses two types of data collection to learn about species distributions: presence/absence and presence-only. The former imagines some version of designed sampling where say plots (grid cells, quadrats, etc.) are sampled and presence/absence or abundance of a species is observed for the sampled plots. That is, locations are fixed. Presence-only data is imagined in terms of randomly encountering a species within a region and is typically collected in the form of museum or citizen science data. That is, locations are random. In fact, the distinction between the two types of data collection can be murky since, if data collection is viewed through gridding of cells, then, conceptually, the observations associated with the cells can be imagined as capturing presence/absence as well as presence-only, as we elaborate below. In any event, the literature on modeling presence/absence data is enormous by now and, more recently, there has been a consequential growth in the literature addressing modeling for the presence-only setting. References to this literature are supplied as part of the development in Sections 3 and 6, respectively.

The contribution of this paper is to address some fundamental and occasionally contentious threads in the literature with regard to the foregoing data collection. For instance, it is asserted that a common modeling framework can be used for both data types, that presence/absence data modeling can be induced under a presence-only framework, and, moreover, that presence-only data can be used to infer about presence/absence (Dorazio, 2014; Royle et al., 2012; Hastie and Fithian, 2013). A further implication is that fusion of general presence/absence and presence-only data sources can be implemented within what is essentially the presence-only framework (Pacifici et al., 2017).

We step into the fray first with discussion to attempt to clarify what “presence at a location” means. We argue that probabilistic modeling for the two data types is distinct and incompatible. Specifically, we argue in detail that presence/absence data should be modeled at point-level. Next, under point-level modeling for such data, we bring in preferential sampling ideas to clarify how potential bias in sampling locations can affect inference with regard to presence/absence. Using the “shared process” perspective, we demonstrate that estimation of the probability of presence as well as prediction of presence can be improved by accounting for preferential sampling. Then, we briefly turn to the fusion problem, again arguing that current versions of such fusion in the literature have fundamental flaws. We propose a probabilistically coherent fusion, again employing the shared process perspective for implementing the fusion, extending application of preferential sampling. This allows the two data sources to be probabilistically independent or dependent. Altogether, this perspective enables a collection of models to take presence/absence modeling to a much richer explanatory level.

In order to examine these contentious issues, we need to spend some time with the presence/absence literature, describing customary modeling. We also need to offer the same with regard to the presence-only literature. Evidently, we also need to elaborate what preferential sampling is in order to reveal its utility for these issues. In the interest of keeping the explication at a concise and comfortable level, we only consider individual species models. However, extension to joint species distribution modeling is available and will be presented in a subsequent paper. In order to go forward, we first need some preliminary words regarding what a presence/absence event means.^1^

### 1.1 The fundamental issue

The fundamental issue that motivates the development of this paper is the attempt to clarify exactly what a presence/absence event means? It seems that if one asks different ecologists one may get different answers. Here, we seek to provide a *coherent* definition, a definition which enables *generative* probabilistic modeling for presence/absence data.

In order to illuminate the issue, we focus on plants (in order to remove movement challenges). Then, for a given species, we can ask what the true realization of a presence/absence surface over a fixed region at a *fixed* time would look like? Evidently, this surface must, at any location in the region, take on the value of either 1 if a presence is there or 0 if not. Critically, this surface can not be imagined through areal units. That is, if it is imagined to be 1 for some units and 0 for other units, then the realization of the surface is clearly dependent upon the arbitrary selection of the units, their size, their shape, their orientation. This would seem to be at odds with how presence/absence arises in nature. In fact, it argues that, if presence/absence is to be viewed coherently, it must be viewed at *point* level. Here, “coherently” means a probabilistically generative specification, a specification for presence/absence that could produce the true realization of the surface. Specifically, from a point level definition, we can scale up to arbitrary areal units (see below) whereas we can not do the reverse.

More explicitly, from a point-level perspective, we can “see” the realization of the presence/absence surface over the entire region of interest, and thus, over any subset of the region. We don’t have to impose any areal scales. In fact, as we argue below, we can attempt to clarify the behavior of this surface. This remark reveals the disconnect in definition in the literature. Presence/absence data, due to the nature of data collection, is frequently associated with areal units, e.g., described as presence/absence over a grid cell, e.g., a plot or a quadrat. The disconnect is exacerbated by scaling. Depending upon the size of the region relative to the size of the areal units, the unit may be considered as a point in the region. However, formally, presence/absence is *never* observed at a point. In practice, a point is only specified with regard to a number of significant decimal places so, implicitly, it is an area due to rounding. The idea of a point-level process specification is accepted as conceptual.^2^

When presence/absence data is referred to at areal units, presence is customarily declared if the species is found anywhere in the unit. But then, it seems impossible to model the probability of presence without considering the size of the unit. Moreover, the definition would ignore the abundance on the unit (suggesting that abundance may be more useful than presence/absence). Should presence associated with one individual in a unit be the same as presence associated with ten individuals in a unit of the same size? Shouldn’t there be implications for probability of presence in the unit? Again, coherence finds us wanting to think of presence/absence in a *unitless* fashion.

It can be argued that the presences of a species over a region form a point pattern. That is, there are a random, finite number of individuals randomly located in the region. We agree and pursue this line of thinking more precisely below. However, the connection to a realization of a presence/absence surface must be made carefully, as well as to a model for a presence/absence probability surface. In this regard, would an ecologist who went out to sample a fixed collection of units for presence/absence attempt to model presence/absence through a (partial) realization of a point pattern? Evidently, the answer is no. Some version of a regression model using suitable unit-level covariates would be attempted, as we elaborate below.

Indeed, under probabilistic thinking, we would imagine a Bernoulli trial at every location in the region. Then, a realization of a presence/absence surface would become a binary map, i.e., a 1 or 0 at each point in the region. This surface is unobservable; it is conceptual and we can think about what it should look like, what properties it should present. In terms of “seeing” the this surface, at best we can display it with a high resolution grid of points. However, to model this surface we need to define a probability of presence at every location, hence, a probability of presence surface over the region. Below, we attempt to clarify more about the behavior of this surface. However, for example, this surface can be thresholded in order to obtain a *niche* for the species within the region.

This also emphasizes that there is no notion of an *intensity* associated with presence/absence observations. Intensities arise from thinking about presence/absence through point patterns, a perspective that is associated with presence-only data, as we develop below. Intensity surfaces can be normalized to density surfaces. Such density surfaces attach probabilities to areas and have nothing to do with a surface of presence probabilities.

Continuing, if we scale a realization of a presence/absence surface to an areal unit then it makes sense to think about the average of the realization over that unit, i.e., the proportion of 1’s over the unit. This proportion is the *empirical* chance of finding a presence at a randomly selected location within the unit. In fact, the proportion of 1’s over the entire region is naturally interpreted as the prevalence of the species over the region. Similarly, with a modeled probability of presence surface, if we scale this surface over the unit, we obtain an average probability over the unit. This average conveys the modeled probability of finding a presence at a randomly selected location within the unit. Again, the issue is that we need not think in terms of areal units in order to model presence/absence. If we want to impose units then we can scale accordingly.

## 2 Our motivating dataset

We illustrate all of the above with invasive plant data from New England in the U.S. We extracted a subregion of the six New England states (Connecticut, Rhode Island, Massachusetts, Vermont, New Hampshire and Maine). The presence/absence dataset comes from the Invasive Plant Atlas of New England (IPANE) and consists of more than 4000 sites where invasive species surveys were conducted and focuses on seven species. Details are provided below. The presence-only dataset comes for the Global Biodiversity Information Facility (GBIF) which is a data aggregator for biological collections worldwide. The number of observations will vary from species to species. Again, details are provided below.

IPANE is a citizen science organization that engages volunteers in scientifically rigorous sampling protocols. There are 4314 unique sampling sites across New England where invasive plant surveys were conducted. Each site is provided with a location (latitude, longitude) and has been classified with regard to each focal species as a presence (focal species recorded) or an absence (focal species not recorded). The dataset includes seven of the most common invasive plant species in the IPANE database: multiflora rose (MR), oriental bittersweet (OB), Japanese barberry (JB), glossy buckthorn (GB), autumn olive (AO), burning bush (BB) and garlic mustard (GM). All species are terrestrial and all but garlic mustard are woody (shrubs, small trees, or vines). These species vary in their land cover associations (e.g., some occur in forest understory and others occur in open habitats). We consider the same species within GBIF. Duplicated points and points lying outside the study region are discarded from the original dataset. Table 1 shows the species name and sample size for the IPANE and the GBIF datasets. In the analysis below, for convenience, longitude and latitude are transformed to eastings and northings, and rescaled from km units to 10 km units.

**Table 1:** Study species and sample sizes

| Common<br>name | symbol | IPANE<br>presences | IPANE<br>absences | GBIF<br>presences |
| --- | --- | --- | --- | --- |
| multiflora rose | MR | 1230 | 3084 | 249 |
| oriental bittersweet | OB | 1106 | 3208 | 305 |
| Japanese barberry | JB | 1012 | 3302 | 399 |
| glossy buckthorn | GB | 755 | 3559 | 223 |
| autumn olive | AO | 386 | 3928 | 193 |
| burning bush | BB | 336 | 3978 | 257 |
| garlic mustard | GM | 279 | 4035 | 440 |

Figure 1 shows the distribution of presence and absence locations from IPANE for each species across the study region. Figure 2 shows the distribution of presence-only points from GBIF for each species across the study region. For some species, for example garlic mustard, the distribution of the presence-only points shows a different pattern from that of the observed presences in the presence/absence data. Again, the presences in IPANE arise from fixed sampling locations while the presences in GBIF arise at random locations. Importantly, we removed all of upper Maine as the figures show. Both the IPANE and the GBIF data were so sparse there that extending spatial modeling to include this region produced poorly behaved model fitting.

**Figure 1:**
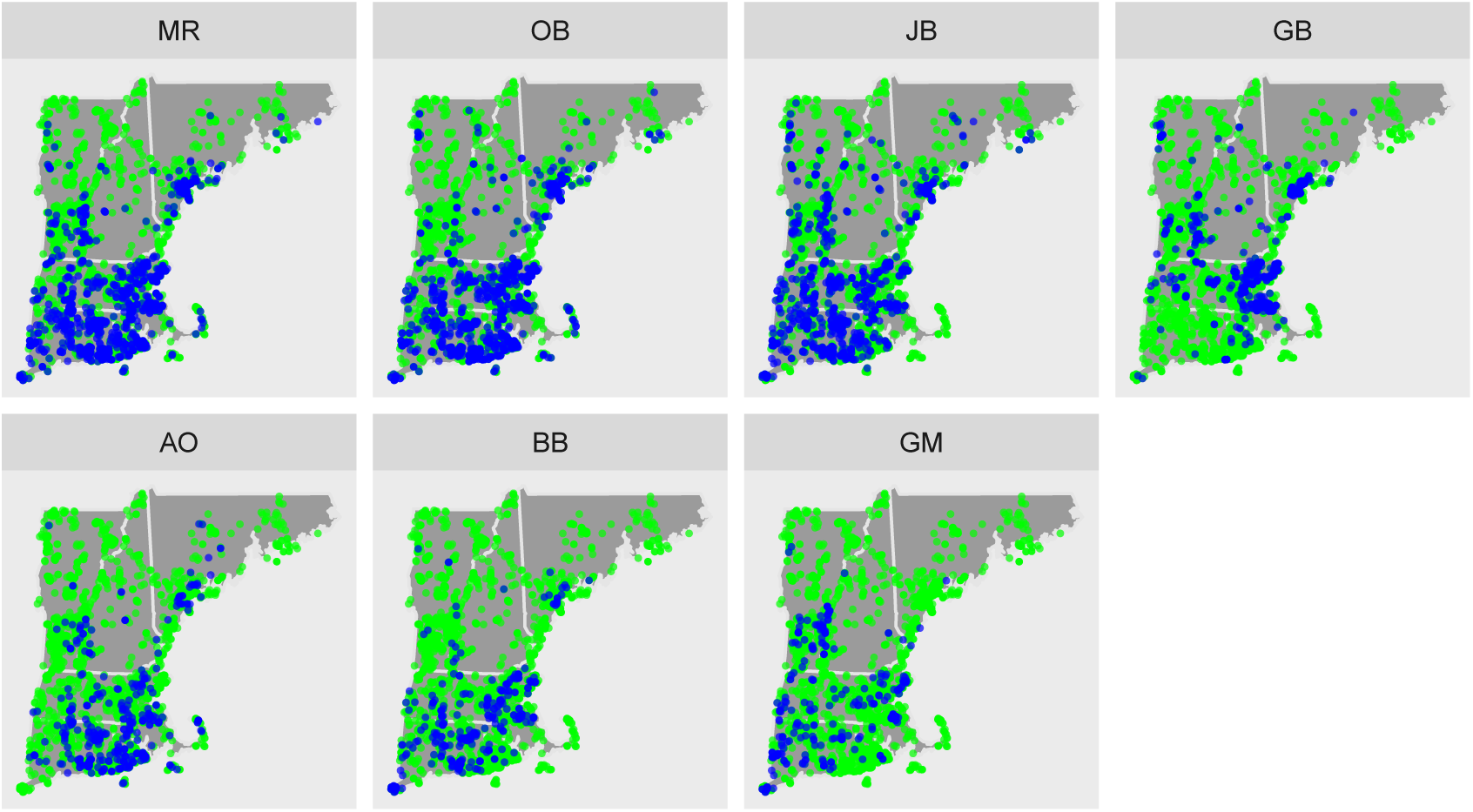
The distribution of presence (blue) and absence (green) points for each species across the study region.

**Figure 2:**
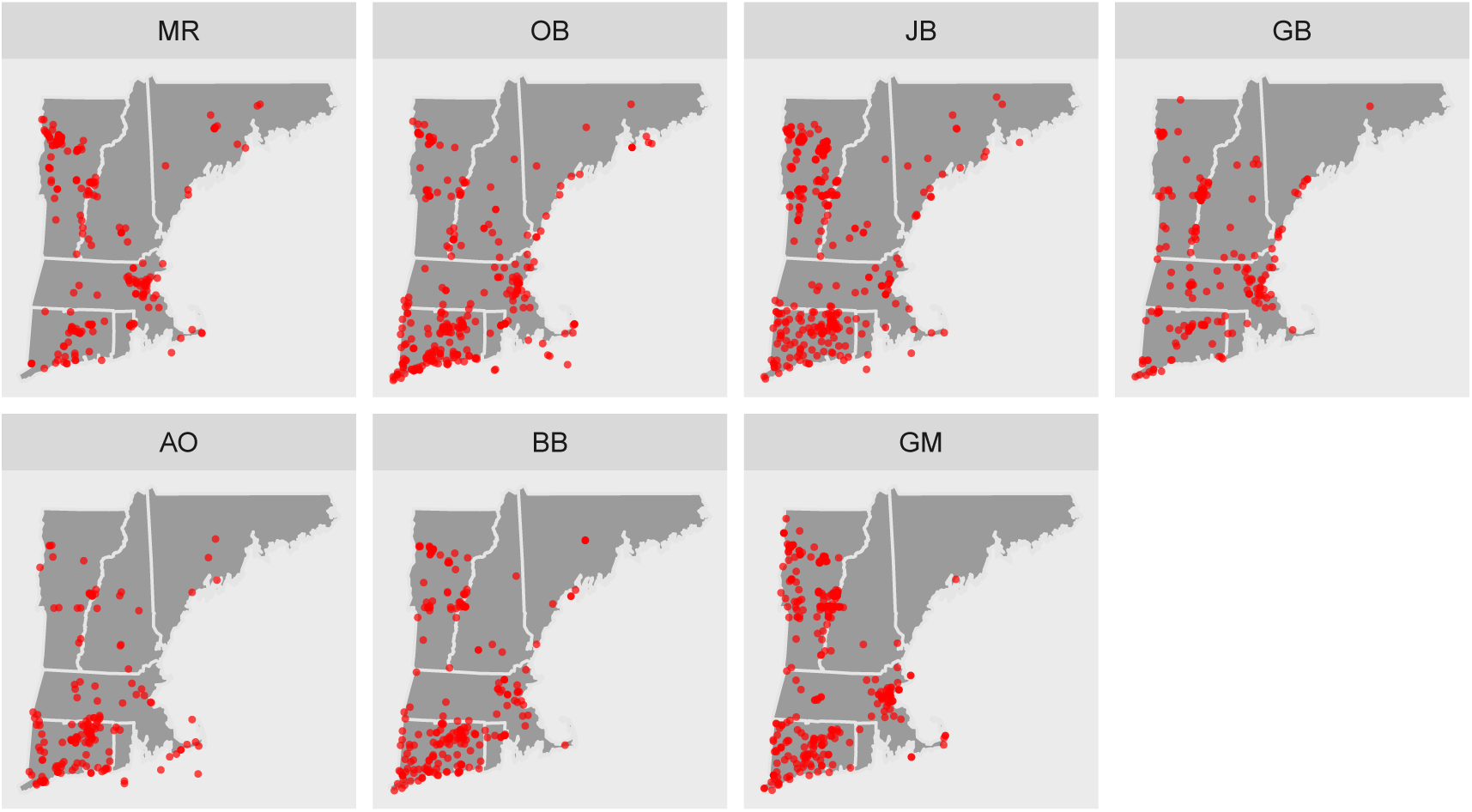
The distribution of presence only points (red) for each species across the study region.

Adding to the original database, we have 19 potential covariates provided by WorldClim (version 1.4, http://www.worldclim.org/version1) as 30-arc second (∼1 km) raster data. We select 7 covariates from them by discarding highly correlated covariates. They are (1) mean diurnal range (mDR, mean of monthly (max temp-min temp)), (2) max temperature of warmest month (max-TWM), (3) min temperature of coldest month (minTCM), (4) mean temperature of driest quarter (meanTDQ), (5) precipitation of wettest month (PWM), (6) precipitation seasonality (PS, the standard deviation of the monthly precipitation estimates expressed as a percentage of the mean of those estimates, that is, the annual mean), and (7) precipitation of warmest quarter (PWQ). These seven covariates were chosen such that each pair has absolute correlation less than 0.7.

## 3 Some presence/absence modeling details

Again, presence/absence data views the observations as binary responses, presence (1) or absence (0) at a collection of sampling locations (see, e.g., Elith et al., 2006, and references therein for a review). The goal is to explain the probability of presence at a location given the environmental conditions that are present there. The natural approach is to build a binary regression model with say a logit or probit link where the covariates can be introduced linearly (see below) or as smoothly varying functions. The latter choice results in generalized additive models (GAMs) which tend to fit data well since they employ additional parameters to enable the response to assume nonlinear and multimodal relationships with the predictors (Guisan et al., 2002; Elith et al., 2006). The price that is paid for using GAMs is a loss of simplicity in interpretation as well as the risk of overfitting with poor out-of-sample prediction. We don’t consider GAMs further here.

Much of the earlier presence/absence work was *non-spatial* in the sense that, though it included spatial covariate information, it did not model anticipated spatial dependence in presence/absence probabilities. Accounting for the latter seems critical. Causal ecological explanations such as localized dispersal as well as omitted (or unobserved) explanatory variables with spatial pattern such as local smoothness of soil or topographic features suggest that, at sufficiently high resolution, occurrence of a species at one location will be associated with its occurrence at neighboring locations (Ver Hoef et al., 2001). In particular, such dependence structure, introduced through spatial random effects, facilitates learning about presence/absence for portions of a study region that have not been sampled, accommodating gaps in sampling and irregular sampling effort.

To begin with some modeling, suppose *Y* (**s**) denotes the presence/absence (1/0) of the species at sample location **s**. If the study region *D* is partitioned into grid cells, say at the level of resolution of the environmental covariates, then, summing up *Y* (**s**) over the number of sample sites in cell *i* (*n*_*i*_) yields grid cell level counts: *Y*_*i*+_ =Σ **s***∈*grid*i Y* (**s**). This is an elementary illustration of scaling up from points to areal units. If the sampling site is viewed as the grid cell then we have *n*_*i*_ =1, a single Bernoulli trial for the cell. If the cell was not sampled, we have *n*_*i*_ =0.

If we assume independence for the trials, a binomial distribution results for *Y*_*i*+_, i.e.,

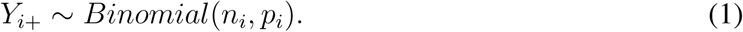

Explicitly, the probability that the species occurs in cell *i, p*_*i*_, is related functionally to the environmental variables with a logit link function and a linear (in coefficients) predictor 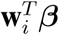, e.g., 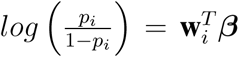. Here **w**_*i*_ is a vector of explanatory environmental variables associated with cell *i* and ***β*** is a vector of associated coefficients. Here, and in the sequel, we could equally well use a probit link function.

If we model probability of presence viewing sites as points, *Y* (**s**) would be taken as

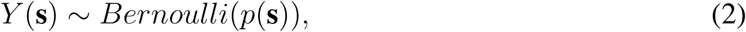

analogously relating the probability that the species occurs in site **s**, *p*(**s**), to the set of environmental variables as 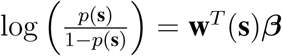. Such modelling requires that we have covariate levels **w**(**s**) for each site. This model is referred to as a spatial regression in the sense that the regressors are spatially referenced. If we set **w**(**s**) =**w**_*i*_ when **s** is within grid *i*, we return to the same model as in (1).

Next, we extend to a simple, grid cell level, spatially explicit model by adding spatial random effects. In modeling *p*_*i*_, a spatial term *ρ*_*i*_ associated with grid *i* is added yielding

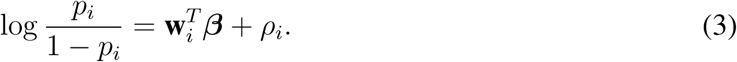

The random effect *ρ*_*i*_ adjusts the probability of presence of the modeled species up or down, depending on the values in a *spatial neighborhood* of cell *i*. To capture this behavior, we customarily employ a Gaussian intrinsic or conditional auto-regressive (CAR) model (Besag, 1974). Such a model proposes that the effect for a particular grid cell should be roughly the average of the effects of its neighboring cells and results in a multivariate normal as the joint distribution over all the cells. There are many ways to specify neighbor structure; see Banerjee et al. (2014) for a full discussion.

Most relevant for us for the remainder of this paper, we confine ourselves to point level spatial model, extending (2). For such data, spatial dependence between points can be modeled based on their relative locations, using Gaussian processes, creating geostatistical models (Banerjee et al., 2014). We would model *Y* (**s**) given *p*(**s**) and augment the explanation of *p*(**s**) through the form

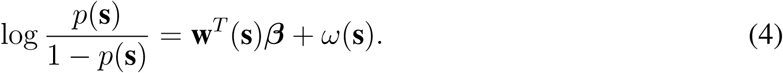

Here, *ω*(**s**) is the spatial random effect associated with point **s**, arising as a realization of a Gaussian process. A suitable covariance function would be selected. With binary response this model is referred to a spatial generalized linear model (GLM); see Diggle et al. (1998). The first stage sampling mechanism is a Bernoulli trial with the surface of probability of presence as a second stage specification. Inference from (4) would be about this surface at any location in the study region, with these probabilities explained through the spatially referenced predictors. Another surface of interest is the realized presence/absence surface, i.e. {*Y* (**s**): **s** *∈ D*}.

The fact that presence/absence is not observable at point level does not preclude useful point level modeling. Indeed, this is the case with all geostatistical modeling (Banerjee et al., 2014), e.g., temperature is never observed at a unitless location but we routinely model temperature surfaces. Taking (4), with say a probit link, *P* (*Y* (**s**) =1) ≡ *p*(**s**) =Φ(**w**^*T*^ (**s**)***β*** + *ω*(**s**)). That is, *P* (*Y* (**s**) =1) =*P* (*Z*(**s**) *>* 0) where *Z*(**s**) =**w**^*T*^ (**s**)***β*** + *ω*(**s**) + *c*(**s**) and *ω*(**s**) is a mean 0 Gaussian process with a suitable correlation function, typically an exponential or, more generally, a Matérn.

Under this model, the *Y* (**s**) are drawn as conditionally independent Bernoulli trials given *p*(**s**) (the *Z*(**s**) are conditionally independent normals). As a result, even if *p*(**s**) is smooth, realizations of the presence/absence surface are everywhere discontinuous. However, below (Section 4) we argue that the realized presence/absence surface should be *locally* continuous. Of course, the *Y* (**s**)’s will be marginally dependent and smoothness of *p*(**s**) will encourage a gridded image of a realization to offer a locally constant (0 or 1) appearance.

An alternative presence/absence specification is a first stage or direct model which introduces a latent Gaussian process at the first modeling stage, setting *Y* (**s**) =1(*Z*(**s**) *>* 0), a function of *Z*(**s**). Now, if *Z*(**s**) is, again, a realization of a Gaussian process which is smooth, then the realized *Y* (**s**) surface will be locally constant. For instance, if *Z*(**s**) =**w**^*T*^ (**s**)***β*** + *ω*(**s**), as above, with an almost everywhere smooth mean surface, we have this behavior. The first stage modeling approach can be attractive for joint species distribution modeling (Clark et al., 2017) since it allows direct modeling of dependence between species rather than deferring it to the second stage (Ovaskainen et al., 2016).

Unfortunately, a technical problem arises in fitting the direct model. This concerns the difference between the probability of presence surface, *p*(**s**), that is, Φ(**w**^*T*^ (**s**)***β*** + *ω*(**s**)) under the second stage model and the realized presence surface under the direct model, 1(**w**^*T*^ (**s**)***β*** + *ω*(**s**) *≥* 0). The realized presence surface has to “agree” well with the observed presences and absences while the probability of presence surface does not. We can observe a presence that has small probability of occurring or an absence that has a small probability of occurring. As a result, the probability of presence surface does not have to work as hard to fit the data. Specifically, with *ω*(**s**) in the modeling, under the direct model, the GP has to react strongly to observed presences and absences. Under second stage modeling, it can react less so. Therefore, when fitting the direct model, the flexibility of the GP results in the *ω*(**s**) surface becoming spiky in the neighborhood of a presence in order to explain well the observed presence.

Can we achieve a locally constant realized presence/absence surface and a smoothed probability of presence surface? A proposal is the following. Still, we let *Y* (**s**) =1, 0 according to *Z*(**s**) *≥* 0, *<* 0. However, we introduce a second GP in specifying *Z*(**s**), i.e., *Z*(**s**) =**w**^*T*^ (**s**)***β*** + *ω*(**s**)+ *γ*(**s**). Here, *ω*(**s**) has a larger range, a smaller decay parameter while *γ*(**s**) has a smaller range with a larger decay parameter. (We are capturing the frequently used interpretation of the “nugget” as microscale dependence (Banerjee et al., 2014)). Then, we define the probability of presence surface as *p*(**s**) =*P* (*Z*(**s**) *≥* 0*|****β***, **w**(**s**), *ω*(**s**)) =Φ(**w**^*T*^ (**s**)***β*** + *ω*(**s**)) while we define the realized presence/absence surface again as 1(*Z*(**s**) *≥* 0). Since *γ*(**s**) is smooth, we will have locally constant behavior in this surface. The *γ*’s will be spiky but the *ω*’s will be smoother. Strong prior information will be needed to control the decay parameters in the GP’s. In fact, we would impose an order restriction on the ranges or decays, demanding more rapid decay for the *γ*(**s**) process. Additionally, we can impose ranges for both the *ω*’s and *γ*’s which are appropriate for the spatial scale of *D* along with the smallest inter-point distance among the presence/absence locations.

If we let *γ*(**s**) be a pure error process, then we would again obtain the problem of *Z*(**s**) being everywhere discontinuous so that the realized *Y* (**s**) surface would be everywhere discontinuous. However, a pure error process with very small variance will provide results similar to that for a GP with very short range, with very rapid decay and the pure error process model will be easier to fit. In fact, in the sequel, we adopt the first stage model with pure error for *γ*(**s**), i.e., *γ*(**s**) ∼ *N* (0, *τ* ^2^) where *τ* ^2^ is fixed for the identifiability of other parameters.

## 4 What does “probability of presence” mean?

We return to the discussion in Section 1.1 regarding what a presence means but now we will be more explicit. Again, the issue is whether presence/absence is viewed at point level or at areal level. Is it a Bernoulli trial at a location or is it the probability that the number of individuals of a species in set *A* is *≥* 1? If we model presence/absence at point level, it is clear what *Y* (**s**) =1 means but what does *Y* (*A*) mean? A coherent probabilistic definition arises as a block average, i.e., a realization of *Y* (*A*) is ∫ _*A*_ 1(*Y* (**s**) =1)*d***s***/|A|* (where *|A|* is the area of *A*), the proportion of the *Y* (**s**) in *A* that equal 1; it is not a Bernoulli trial and *P* (*Y* (*A*) =1) =0. We can calculate *E*(*Y* (*A*)) =∫_*A*_ *p*(**s**)*d***s***/|A|* with *p*(**s**) as in (4). That is, *E*(*Y* (*A*)) becomes the average probability of presence over *A*. It is the probability that the species is present at a randomly selected location in *A*.

If *p*(**s**) is constant over *A* then *E*(*Y* (*A*)) is this constant probability. This takes us back to the case of gridded regions where we defined *p*_*i*_, the constant probability over *A*_*i*_ using logistic (or probit) regressions, as in the previous section. Importantly, that areal definition of *p*_*i*_ is interpreted at point level; it is the probability of presence at any site in *A*_*i*_.

Now, suppose we consider the locations of all individuals in a study region as a random point pattern. Then, if *N* (*A*) is the number of individuals in set *A, P* (presence *in A*) =*P* (*N* (*A*) *≥* 1). Here, assuming a nonhomogeneous Poisson process or, more generally a log Gaussian Cox process (LGCP) with intensity *λ*(**s**) (see Section 5.2 below), *N* (*A*) ∼ *Po*(*λ*(*A*)) where *λ*(*A*) = ∫_*A*_*λ*(*s*)*ds*. Then *P* (*Y* (*A*) =1) =*P* (*N* (*A*) *≥* 1) =1 *-e*^*-λ*(*A*)^. Since presence-only data alleges to sample the point pattern (although likely not fully but, rather, up to sampling effort over the region (Chakraborty et al., 2011; Fithian et al., 2015), it is compatible with this definition of presence/absence. However, the occurrence probability is only defined with regard to the size of *A*, a concern raised in Hastie and Fithian (2013); evidently, occurrence probability will vary with the size of *A*. As a result, it is unclear how to specify a meaningful probability of presence surface. Furthermore, the definition of probability of presence as “one or more” observations of the species in *A* yields local distortion to any such surface; *N* (*A*) =1 and *N* (*B*) =11 are treated the same with regard to presence if *|A|* =*|B|* (Aarts et al., 2012). Moreover, even if we ignore the size of *A* and return to a grid of cells over *D*, then it is clear that *p*_*i*_ ≡ *P* (*Y* (*A*_*i*_) =1) has nothing to do with *p*_*i*_ defined in the previous section.

The two foregoing definitions associated with *P* (presence *in A*) are incompatible and the fundamental difference between them has been ignored in the literature. The conceptualization for the first choice is that we go to fixed “point” locations and see what is there; we are not sampling a point pattern. There is a surface over a domain *D* which captures the probability of presence at every location in *D*. The conceptualization for the second is that we identify an area of interest *D* and we census it completely for all of the occurrences of the point pattern in it. It provides an intensity surface which can be scaled to a density surface. However, as with any probability density function, the density surface at a point is not the probability of presence at that point. The second version can not scale down to point level since then *λ*(*A*) *→* 0.

Furthermore, if presented with a collection of plots and observed presence/absence for those plots, would one ever model the data as a point pattern? The answer seems clear; no point pattern was observed, there is no way to model an intensity. We would use one of the foregoing presence/absence models. Moreover, if we briefly consider the data fusion problem, suppose one obtains an additional set of presence-only data for the region. Why is it now appropriate to model the original presence/absence data using a point pattern model associated with the presence-only data?

So, we have articulated the issue with regard to being too informal in terms of the notion of presence as well as the data fusion challenge and have asserted that modeling presence/absence at the point level seems the preferable specification. Of course, one can disregard the scaling issue, create an arbitrary discretization of the space, and calculate probabilities over the discretization, as in recent work of Pacifici et al. (2017). Furthermore, in the literature to date, ignoring the incompatibility is the way that presence-only data has been used to provide presence/absence probabilities and also the way presence-only data has been fused with presence/absence data. We propose remediation for this in Sections 5 and 6 below but first, in Section 4, we attempt further clarification of point-level presence/absence modeling.

We briefly digress to a related contentious issue in the recent literature. Can one use presence-only data to infer about presence/absence? Fundamentally, the answer must be no since one needs to see absences in order to model probability of presence (see Section 6.1 below in this regard). However, Royle et al. (2012), imagining an areal unit definition of presence, argue that “occurrence probability can be estimated from presence-only data.” In particular, they assume that an environmental covariate, *z*, is a priori, uniformly distributed over the study region. Then, with *P* (*Y* (*A*) =1*|z*(*A*)) =*ψ*(*β*_0_ + *β*_1_*z*(*A*)) for link function *ψ* and a discrete uniform density for *z*(*A*), using Bayes’ Theorem,

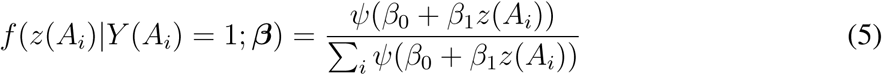

Equation (5) suggests that, by modeling environment/habitat given presence, we can learn about *P* (*Y* (*A*_*i*_) =1*|z*(*A*_*i*_)). Hastie and Fithian (2013) point out that this model is flawed in the sense that the unconditional probabilities, *P* (*Y* (*A*_*i*_) =1) are not identified; only relative probabilities are identifiable. We add two further comments. First, the likelihood associated with (5), in fact, is Π_*i*_*ψ*(*β*_0_ + *β*_1_*z*(*A*_*i*_))*/c*(*β*_0_, *β*_1_). This is a different function of the *β*s than the likelihood for a binary regression with *P* (*Y* (*A*_*i*_) =1*|β*_0_, *β*_1_, *z*(*A*_*i*_)). The parameters do not mean the same thing in the two models and would not provide the same estimates if we could fit the latter. Second, from above, it is unclear what the event *Y* (*A*_*i*_) =1 means and, regardless, the occurrence probabilities being considered here suffer the same issues as above with regard to the size of the *A*_*i*_.

### 4.1 Further clarification of point level presence/absence modeling

A critical issue in reconciling the differences above is to think more carefully about what the distribution of a species looks like within a specified region, *D*. Suppose we consider the complete census of individuals in the region. To be realistic, we have to view the number of presences in a bounded region as finite and therefore a presence must be bigger than a (unitless) point since there are an uncountable number of points in *D*. Again, the scaling issue arises. Formally, a presence can not arise at a point, it is not unitless in size; practically, it can be point-referenced. That is, formally, the presence/absence surface over the region consists of a finite set of “patches” where the species is present and, outside of these patches, the species is absent. From an ecological and practical perspective, we could think of a patch as a collection of individuals of a particular species (it might be just one) that is dense enough so that, at point level, we would declare presence for every location in the patch. However, if the gaps between the individuals become sufficiently large, then those locations in the gaps must now become absences. The scaling here is qualitative, not quantitative - an ecologist would not attempt to be precise here and the denseness needed to define a patch evidently depends upon the sizes of the patch relative to the size of *D*. In the sequel, we also avoid defining patch sizes.

Then, a presence-only realization becomes this finite set of patches. To view it as a point pattern, we might assume the patches are small regions about the individuals and the observed point is say, the centroid of its patch. To make this definition consistent, we have to assume that only one point is associated with a patch. That is, with a complete census, the number of patches equals the number of points in the point pattern. However, with regard to presence/absence, a presence at a location is observed if the location falls within a patch associated with a point. This definition of the realized presence/absence surface gives an immediately rigorous definition of prevalence. The prevalence of the species over *D* is the total area of the patches for the species relative to the total area of *D*.

Consider some important implications. First, the number of presence points in *D* is uncountable, as is the number of absence points. Second, presence/absence is a *neighborhood* phenomenon. If there is a presence at **s** then there is presence everywhere in a sufficiently small neighborhood, *∂***s**, of **s**. Similarly, if there is an absence at **s**, then there must be a neighborhood of **s** where every location is an absence. As a result, the *realized* presence absence surface is locally constant. A suitable probability model for presence/absence should provide realizations which are locally constant. This returns us to the discussion at the end of Section 3. A model which assumes conditionally independent Bernoulli trials across locations is not formally appropriate since such a model will provide random 0s and 1s across locations, yielding no local smoothness.

Third, conceptually, the number points in the point pattern can be smaller or larger than the number of observed presences. That is, observing a presence at a location is not identical to observing the centroid associated with the patch containing the observed presence. According to selection of sampling sites, the same individual may be observed at more than one point (though, in practice, it is not likely to be recorded as such) but also, some individuals may never be observed. Practically, we acknowledge that presence/absence sampling will never observe all individuals but also, that presence-only sampling will rarely observe all individuals. So, with a dataset of point-level presence/absence locations and a dataset of presence-only random locations, at sufficiently high spatial resolution, the two sets of locations will be disjoint.^3^

## 5 Preferential sampling

We propose preferential sampling as a tool for both improving presence/absence prediction as well as for fusing presence-only data with presence/absence data. To begin we need to formally develop the concept of preferential sampling.

### 5.1 What is preferential sampling all about?

The notion of preferential sampling was introduced into the literature in the seminal paper of Diggle et al. (2010). Subsequently, there has been considerable follow up research. Two useful papers in this regard are Pati et al. (2011) and Cecconi et al. (2016). A standard illustration arises in geostatistical modeling (see e.g. Cressie and Wikle, 2011; Banerjee et al., 2014). Consider the objective of inferring about environmental exposures. If environmental monitors are only placed in locations where environmental levels tend to be high, then interpolation based upon observations from these stations will necessarily produce only high predictions. The obvious remedy lies in suitable spatial design of the locations, e.g., a random or space-filling design (Saltzman and Nychka, 1998) for locations over the region of interest is expected to preclude such bias. However, sampling for presence/absence may not be designed in this fashion; ecologists may tend to sample where they expect to find individuals, introducing *bias* into the collection of sampling locations. Recognizing the possibility of such bias, can we revise presence/absence prediction to adjust for it? This is the intention of preferential sampling modeling.

We proceed as follows. While the set of sampling locations may not have been developed randomly, we study it as if it were a realization of a spatial point process. That is, it may be designed in some fashion and be deterministic but not necessarily with the intention of being roughly uniformly distributed over *D*. Then, the question becomes a stochastic one: is the realization of the responses independent of the realization of locations? If no, then we have what is called preferential sampling. Importantly, the dependence here is stochastic dependence. Notationally/functionally, the responses are associated with the locations. We will make this more clear below.

In our context, the presence/absence data has an associated probability of presence surface, as we develop below. This surface plays the role of the “exposure” surface, with the finite set of binary responses, *𝒴*, informing about it. Taking the set of sampling locations as a realization of a random point pattern, *𝒮*, the question we ask is whether *𝒴* is independent of *𝒮*, again in a stochastic sense? Below, we develop several models, using the idea of a *shared* process, that enable us to address this question and, furthermore, whether *𝒮* enables us to improve our inference regarding the presence/absence surface, our prediction of presence.

### 5.2 Preferential sampling models for presence/absence data

To develop the stochastic specifications that formalize preferential sampling for a region *D*, we imagine two cases for the intensity associated with the point pattern of sampling locations, *𝒮*:

i. *λ*(**s**) =**w**^*T*^ (**s**)***β***, i.e., a nonhomogeneous Poisson process (NHPP) and
ii. *λ*(**s**) =**w**^*T*^ (**s**)***β*** + *η*(**s**), a logGaussian Cox process (LGCP).

Here, **w**(**s**) is a vector of predictors with associated regression coefficients ***β*** and *η*(**s**) is a mean 0 GP with a suitable covariance function. See, e.g., Illian et al. (2008) for full discussion of NHPPs and LGCPs.

Consider modeling for *𝒴*. Since we model *Y* (**s**) directly through a latent Gaussian process, *Z*(**s**), i.e., *Y* (**s**) =1(*Z*(**s**) *>* 0), as at the end of Section 3, we only need to propose models for *Z*(**s**).^4^ We start with a simple spatial regression,

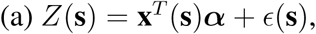

where the predictors in **x**(**s**) and those in **w**(**s**) need not be identical. Extension to a customary geostatistical model for *Z*(**s**) (Banerjee et al., 2014) becomes

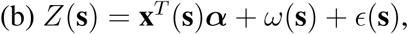

adding *ω*(**s**) as a mean 0 GP, independent of *η*(**s**) above.

To illuminate the model structure, denote the point pattern over *D* by *𝒮*, the realization of *ω* over *D* as ***ω***_*D*_, and the realization of *η* over *D* as ***η***_*D*_. Suppose we consider the joint distribution [*𝒮, 𝒴*, ***ω***_*D*_]. We have the natural factorization as [***ω***_*D*_][*𝒮|****ω***_*D*_][*𝒴| 𝒮*, ***ω***_*D*_] (suppressing ***η***_*D*_ if case (ii)). Then, we say that there is no preferential sampling if [*𝒮|****ω***_*D*_] =[*𝒮*]. This is clearly the case with model (a) or (b) inducing *𝒴* and (i) or (ii) for *𝒮*. Only ***ω*** _*𝒴*_ ={*ω*(**s**_*i*_): **s**_*i*_ ∈ *𝒮*} is needed to fit (see the Supplementary Material).

Now, we can extend model (i) to

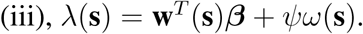

In this notation, with model (b) for *𝒴, ω*(**s**) is a shared process for both *𝒴* and *𝒮* so *𝒴* and *𝒮* are not independent. Working with (b) and (iii), if *ψ* =0, then, following Diggle et al. (2010), we have non-preferential sampling while if *ψ* ≠ 0, we have *strong* preferential sampling.

Pati et al. (2011) extended this idea so that *𝒴* follows the geostatistical model (b) while *𝒮* follows model (ii). Then, they attempt to interpret *η*(**s**) as a regressor to add to the geostatistical model for *𝒴*. That is, now we have model

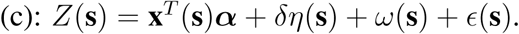

Here, the coefficient *δ* plays a preferential sampling role. For example, suppose the design *𝒮* over-samples locations in *D* where we have presences, where *Y* (**s**) tends to be 1, i.e., where *Z*(**s**) tends to be high. Then, *η*(**s**) will tend to be high around those locations. Therefore, *η*(**s**) can be a significant predictor for *Z*(**s**) (hence for *Y* (**s**)) with *δ >* 0. (A similar argument applies when *δ <* 0.) With (ii) and (c), *η*(**s**) is the shared process. Only ***η***_*𝒴*_ ={*η*(**s**_*i*_): **s**_*i*_ *∈* 𝒮 *}* is needed to fit (see the Supplementary Material).

A further shared process model for *𝒴* that can be explored in this regard extends (a) to

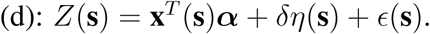

Here, interest is in comparing (d) and (ii) with (a) and (ii); is *δ* ≠ 0? Diggle et al. (2010) focus on comparing (b) and (i) with (b) and (iii). Pati et al. (2011) focus on comparing (ii) and (b) with (ii) and (c).

Cecconi et al. (2016) add another GP to the intensity for 𝒮, i.e.,

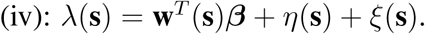

That is, using model (iv) with model (c), there is a shared GP for *𝒴* and *𝒮* as well as individual GP’s for each, a total of three independent GP’s altogether. They acknowledge identifiability problems in model fitting with the three latent Gaussian fields.

We offer Table 2 which provides a summary of the modeling choices for *𝒴* and *𝒮*. However, in the next subsection, we examine just a subset of possible model comparison. We compare (a) and (ii) with (d) and (ii). That is, [*𝒴| 𝒮*, ***α***][*𝒮|****β***, ***η***_*D*_] vs. [*𝒴| 𝒮*, ***α***, ***η***_*𝒴*_, *δ*][*𝒮|****β***, ***η***_*D*_]. We compare (b) and (ii) with (c) and (ii). That is, [*𝒴| 𝒮*, ***α***, ***ω***_*𝒴*_] [*𝒮|****β***, ***η***_*D*_] vs. [*𝒴| 𝒮*, ***α***, ***ω***_*𝒴*_, ***η***_*𝒴*_, *δ*][*𝒮|****β***, ***η***_*D*_]. Model fitting details are given in the Supplementary Material. Since the intent is to improve the predictive performance of the model for *𝒴*, model comparison criteria focuses on out-of-sample prediction for *Y* (**s**)’s. (See Section 5.3)

**Table 2:** Models

| Modeling for $\mathcal{S}$ | | Modeling for $\mathcal{Y}$ | |
| --- | --- | --- | --- |
| (i) | $\lambda(\mathbf{s}) = \mathbf{w}^T(\mathbf{s})\boldsymbol{\beta}$ | (a) | $Z(\mathbf{s}) = \mathbf{x}^T(\mathbf{s})\boldsymbol{\alpha} + \epsilon(\mathbf{s})$ |
| (ii) | $\lambda(\mathbf{s}) = \mathbf{w}^T(\mathbf{s})\boldsymbol{\beta} + \eta(\mathbf{s})$ | (b) | $Z(\mathbf{s}) = \mathbf{x}^T(\mathbf{s})\boldsymbol{\alpha} + \omega(\mathbf{s}) + \epsilon(\mathbf{s})$ |
| (iii) | $\lambda(\mathbf{s}) = \mathbf{w}^T(\mathbf{s})\boldsymbol{\beta} + \psi\omega(\mathbf{s})$ | (c) | $Z(\mathbf{s}) = \mathbf{x}^T(\mathbf{s})\boldsymbol{\alpha} + \delta\eta(\mathbf{s}) + \omega(\mathbf{s}) + \epsilon(\mathbf{s})$ |
| (iv) | $\lambda(\mathbf{s}) = \mathbf{w}^T(\mathbf{s})\boldsymbol{\beta} + \eta(\mathbf{s}) + \xi(\mathbf{s})$ | (d) | $Z(\mathbf{s}) = \mathbf{x}^T(\mathbf{s})\boldsymbol{\alpha} + \delta\eta(\mathbf{s}) + \epsilon(\mathbf{s})$ |

### 5.3 Model fitting and inference for presence/absence data using preferential sampling

To make the model comparison between (b) with (ii) vs. (c) with (ii) we only need to fit the latter and look at the posterior distribution for *δ*. (In fact, since (b) and (ii) are independent, for presence/absence prediction, we only need fit (b).) Similarly, for the model comparison between (a) with (ii) vs. (d) with (ii). (Again, (a) and (ii) are independent.) We do this below for each of the seven species.

We estimate models (a) -(d) for the presence/absence data. For model (c) and (d), we include log Gaussian Cox process models for *𝒮*, i.e., for model (ii), by taking 2,666 regular grid cells over *D* to approximate the region. The regular grid is needed because we introduce the ***η*** surface into models (c) and (d). Among these grid cells, 1870 don’t include any presence/absence locations. The area for each grid cell is standardized as 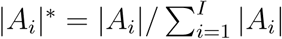 for *i* =1, 2, *…, I* where *|A*_*i*_*|* is the area for grid cell *A*_*i*_. For all species, we use the same seven covariates presented in Section 2 for both **w** and **x**.

As for Bayesian inference, although Gibbs sampling is available for ***ω***, its computational cost/time is *𝒪* (*n*^3^) and required memory is *𝒪* (*n*^2^). In our case, we have a relatively large *n* =4314, so we implement a nearest neighbor Gaussian process (Datta et al., 2016), which is a sparse Gaussian process model whose computational time is *𝒪* (*nk*^3^) (linear in *n*) and required memory is *𝒪* (*nk*) where *k* is the number of neighbors. We set *k* =15 for the analysis below. For sampling ***η***, we implement Metropolis-Hastings (MH) updates. The sampling details for all parameters are described in the Supplementary Material. As for prior specifications, all are weak; we assume 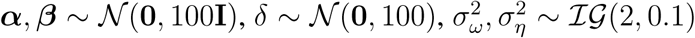 and *ϕ*_*ω*_, *ϕ*_*η*_ ∼*𝒰* (0, 200). We set *τ* ^2^ =1 for the identifiability of other parameters. We discard the first 20,000 iterations as burn-in and preserve the subsequent 20,000 as posterior samples.

Table 3 shows the estimation results for *δ* for models (c) and (d). For MR, JB, GB and AO, the results for model (d) suggest significant preferential sampling effects; the means for *δ* are well away from 0. Again, when *δ >* 0, this implies that, in the selection of the presence/absence locations for the species, presences were oversampled. When *δ <* 0, this means that in the selection of the presence/absence locations for the species, presences were undersampled. This insight is useful but, furthermore, failing to include the *η*(**s**) into the modeling might lead to misinterpretation of the covariate effects.

**Table 3:**
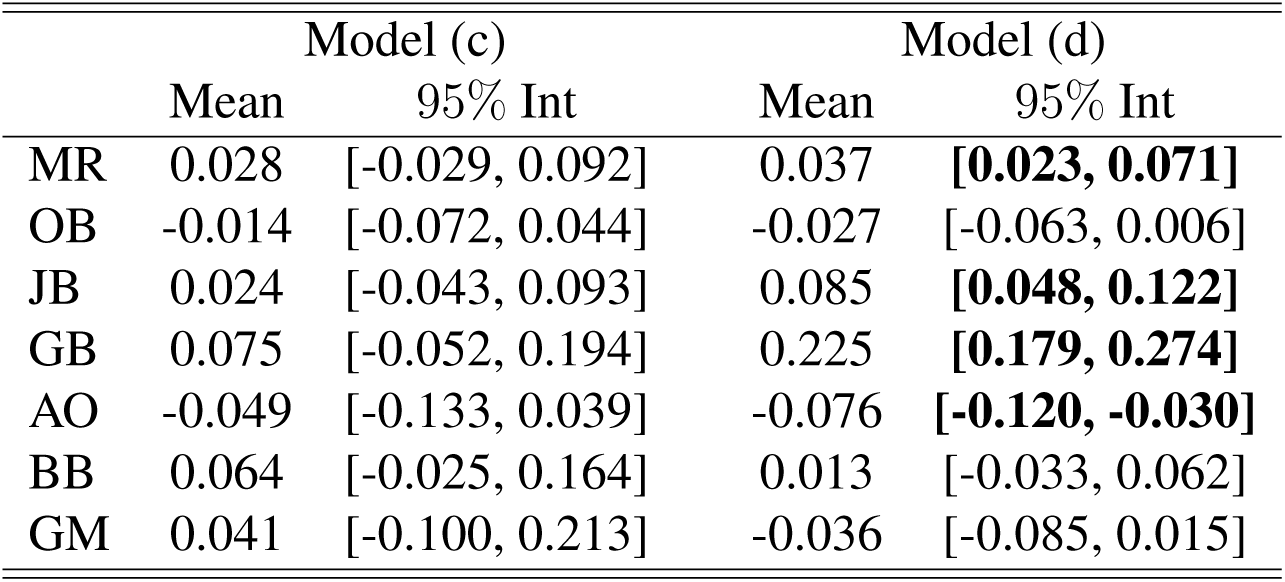
Estimation results for *δ* with models (c) and (d)

|  | Model (c) |  | Model (d) |  |
| --- | --- | --- | --- | --- |
|  | Mean | 95% Int | Mean | 95% Int |
| MR | 0.028 | [-0.029, 0.092] | 0.037 | **[0.023, 0.071]** |
| OB | -0.014 | [-0.072, 0.044] | -0.027 | [-0.063, 0.006] |
| JB | 0.024 | [-0.043, 0.093] | 0.085 | **[0.048, 0.122]** |
| GB | 0.075 | [-0.052, 0.194] | 0.225 | **[0.179, 0.274]** |
| AO | -0.049 | [-0.133, 0.039] | -0.076 | **[-0.120, -0.030]** |
| BB | 0.064 | [-0.025, 0.164] | 0.013 | [-0.033, 0.062] |
| GM | 0.041 | [-0.100, 0.213] | -0.036 | [-0.085, 0.015] |

The *η*(**s**) also provide improved prediction of presence/absence (see below). However, with inclusion of the ***ω*** surface (model (c)), these coefficients become insignificant. Table 4 shows the estimation results for models (a) and (d) for ***α*** for MR, JB, GB and AO, each of whose *δ* is significant with model (d). Introducing the ***η*** surface affects the estimation results for ***α***. For example, the estimated *α* of meanTDQ for GB is significantly negative for model (a) but becomes insignificant for model (d).

**Table 4:** Estimation results for MR, JB, GB, and AO for models (a) and (d). The bold font suggests the change of significance.

|  | MR |  | JB |  | GB |  | AO |  |
| --- | --- | --- | --- | --- | --- | --- | --- | --- |
| Model(a) | Mean | 95% Int | Mean | 95% Int | Mean | 95% Int | Mean | 95% Int |
| const | -1.774 | [-1.951, -1.602] | -1.507 | [-1.676, -1.344] | -1.906 | [-2.126, -1.693] | -2.360 | [-2.634, -2.102] |
| mDR | 0.193 | [0.027, 0.362] | 0.383 | [0.213, 0.556] | 0.110 | [-0.082, 0.305] | 0.381 | [0.161, 0.603] |
| maxTWM | -0.020 | [-0.217, 0.175] | -0.236 | [-0.435, -0.035] | 0.463 | [0.223, 0.704] | -0.554 | [-0.813, -0.300] |
| meanTDQ | 0.715 | [0.460, 0.970] | 0.762 | [0.498, 1.028] | -0.324 | [-0.619, -0.025] | 0.859 | [0.532, 1.196] |
| minTCM | -0.070 | [-0.149, 0.010] | -0.065 | [-0.146, 0.015] | 0.415 | [0.309, 0.522] | 0.029 | [-0.074, 0.136] |
| PWM | 0.178 | [0.092, 0.262] | 0.063 | [-0.024, 0.151] | -0.139 | [-0.240, -0.038] | -0.182 | [-0.300, -0.066] |
| PS | -0.142 | [-0.246, -0.040] | -0.083 | [-0.182, 0.017] | -0.030 | [-0.141, 0.081] | -0.162 | [-0.320, -0.010] |
| PWQ | -0.062 | [-0.166, 0.042] | 0.204 | [0.103, 0.307] | -0.371 | [-0.502, -0.239] | -0.156 | [-0.308, -0.001] |
| Model(d) | Mean | 95% Int | Mean | 95% Int | Mean | 95% Int | Mean | 95% Int |
| const | -1.931 | [-2.156, -1.705] | -1.845 | [-2.072, -1.620] | -2.985 | [-3.331, -2.654] | -2.066 | [-2.391, -1.751] |
| mDR | 0.215 | [0.043, 0.386] | 0.423 | [0.246, 0.597] | **0.352** | **[0.124, 0.578]** | 0.362 | [0.141, 0.585] |
| maxTWM | -0.049 | [-0.247, 0.145] | -0.308 | [-0.514, -0.102] | **0.149** | **[-0.135, 0.433]** | -0.507 | [-0.760, -0.252] |
| meanTDQ | 0.792 | [0.524, 1.054] | 0.900 | [0.622, 1.174] | **0.206** | **[-0.145, 0.550]** | 0.756 | [0.418, 1.089] |
| minTCM | -0.063 | [-0.146, 0.021] | -0.051 | [-0.137, 0.034] | 0.480 | [0.350, 0.612] | 0.020 | [-0.087, 0.129] |
| PWM | 0.161 | [0.073, 0.250] | 0.022 | [-0.070, 0.114] | -0.225 | [-0.371, -0.096] | -0.148 | [-0.269, -0.026] |
| PS | -0.132 | [-0.238, -0.026] | -0.061 | [-0.165, 0.040] | 0.046 | [-0.083, 0.183] | -0.174 | [-0.333, -0.017] |
| PWQ | -0.028 | [-0.141, 0.083] | 0.267 | [0.153, 0.381] | -0.218 | [-0.399, -0.032] | -0.195 | [-0.357, -0.038] |
| $\delta$ | 0.037 | [0.023, 0.071] | 0.085 | [0.048, 0.122] | 0.225 | [0.179, 0.274] | -0.076 | [-0.120, -0.030] |

Figure 3 shows the posterior mean probability of presence surface with models (a) and (c) for JB and GB. The surfaces for model (c) are very different from those for model (a), capturing local behavior. That is, by comparison with Figure 1, model (c) captures the presence probability better than model (a) which smooths away too much detail. This point is supported through comparison of predictive performance below.

**Figure 3:**
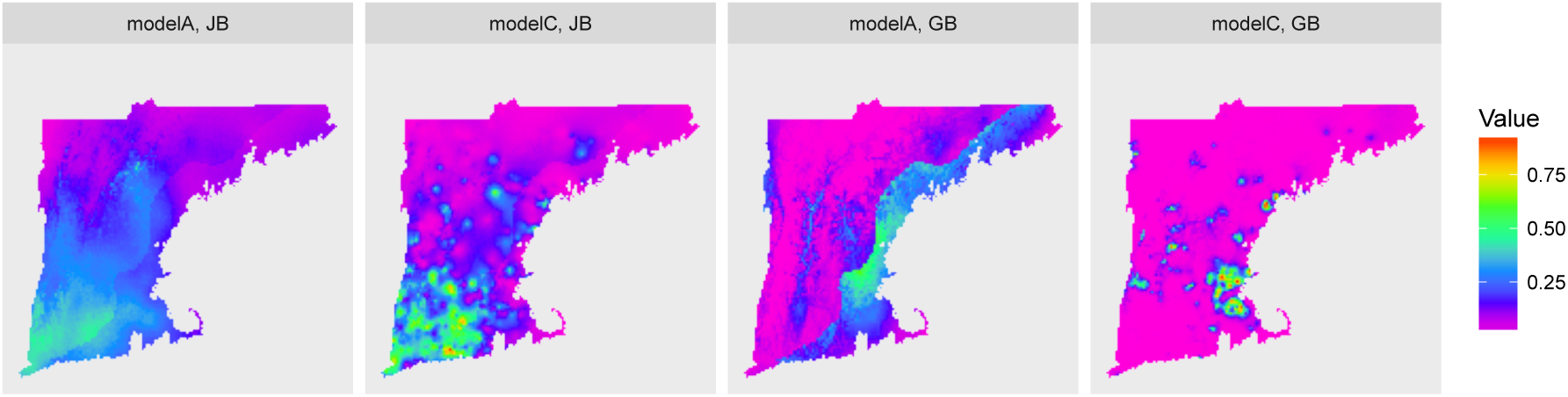
The posterior mean presence probability surface for models (a) and (c) for JB and GB.

To demonstrate improved prediction, with binary response we consider misclassification error using the Tjur *R*^2^ coefficient of determination (Tjur, 2009). This measure prefers a model with high probability of presence when presence is observed and low probability of presence when absence is observed. For species *j*, this quantity is given by 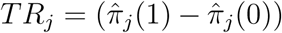 where 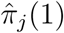 and 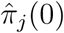 are the average probabilities of presence for the observed ones and zeros associated with the *j*-th species across the locations. The larger the *TR*_*j*_, the better the discrimination. We held out 20% (879) of the PA locations for the seven species. Table 5 shows the results for the TR measure for models (a) -(d). For all species, models including *ω*(**s**), (b) and (c), outperform those without, (a) and (d). Model (c) with (ii) tends to be a better than model (b) (which ignores (ii)) particularly for AO, BB, and GM. Model (d) with (ii) is at least as good as model (a) (which ignores (ii)) but is really only consequentially better for species GB which has the largest *δ* coefficient under model (d).

**Table 5:** Estimation results for the TR measure for preferential sampling.

|  | Model (a) |  | Model (b) |  | Model (c) |  | Model (d) |  |
| --- | --- | --- | --- | --- | --- | --- | --- | --- |
|  | Mean | 95% Int | Mean | 95% Int | Mean | 95% Int | Mean | 95% Int |
| MR | 0.104 | [0.094, 0.114] | 0.168 | [0.145, 0.191] | **0.176** | [0.157, 0.202] | 0.105 | [0.096, 0.114] |
| OB | 0.099 | [0.088, 0.109] | **0.183** | [0.163, 0.200] | **0.183** | [0.158, 0.203] | 0.101 | [0.092, 0.111] |
| JB | 0.072 | [0.063, 0.081] | **0.201** | [0.180, 0.230] | 0.198 | [0.169, 0.227] | 0.075 | [0.066, 0.083] |
| GB | 0.126 | [0.112, 0.139] | 0.405 | [0.372, 0.434] | **0.412** | [0.382, 0.451] | 0.162 | [0.147, 0.175] |
| AO | 0.034 | [0.027, 0.043] | 0.095 | [0.068, 0.129] | **0.115** | [0.073, 0.156] | 0.039 | [0.033, 0.051] |
| BB | 0.057 | [0.048, 0.068] | 0.112 | [0.078, 0.152] | **0.131** | [0.097, 0.168] | 0.057 | [0.049, 0.066] |
| GM | 0.026 | [0.020, 0.032] | 0.135 | [0.111, 0.166] | **0.164** | [0.118, 0.206] | 0.027 | [0.022, 0.033] |

## 6 Fusing presence/absence and presence-only data

We turn to the data fusion question. Data fusion (also *assimilation*) is a widely employed objective when multiple data sources are available to inform about the same response of interest (Nychka and Anderson, 2010; Wikle and Berliner, 2007). A canonical example is the goal of modeling exposure to an environmental contaminant when we might have station data available, computer model output available, and perhaps satellite data as well. The conceptual modeling strategy is to imagine a latent *true* exposure surface and then build a model for each data source, conditioned upon the true surface. The joint modeling enables each of the sources to inform about the true exposure surface, to enable improved prediction of this surface. Examples in the literature include application to weather data, sea surface temperature, and animal behavior patterns (Wikle et al., 2001; Sahu et al., 2016; Rundel et al., 2015).

In our setting, data fusion is different from customary settings. Rather than multiple data sources informing about a common response, e.g., ozone level, we have two different types of data. While both inform about species distribution, we have argued above that presence/absence data is not described stochastically in the same way as presence-only data. The fusion approaches considered in the literature (Fithian et al., 2015; Dorazio, 2014; Giraud et al., 2016; Pacifici et al., 2017) ignore this and assume a latent point pattern model for the presence-only data and that the presence/absence data is induced under this model, as we described above. Since we argue that a point pattern specification is inappropriate for presence/absence data, a different type of fusion is required. We have a point pattern model for the presence-only data and a binary map model for the presence/absence data. So, we again turn to preferential sampling ideas (Diggle et al., 2010) in order to explore a coherent probabilistic fusion.

The extra information available to make a data fusion story is *𝒮* _*PO*_, the set of observed presence-only locations. Formally, what information does *𝒮* _*PO*_ bring with regard to learning about the probability of presence surface? Suppose, as in Section 4, we assume that *𝒮* _*PO*_ is a complete census, i.e., arising from a finite number of individuals which are imagined as blobs in *D* and *𝒮* _*PO*_ are the centroids associated with these blobs. Associated with 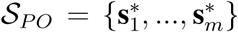, we can imagine a *λ*_*PO*_(**s**) specified with a set of models similar to (i)-(iv). We expect *λ*_*PO*_(**s**) to be elevated near these observations. For example, analogous to (ii), let *λ*_*PO*_(**s**) =**w**^*T*^ (**s**)***β***_*PO*_ + *η*_*PO*_(**s**), using the same predictors as with the presence/absence modeling. Because the mechanisms that created *𝒮* _*PO*_ and *𝒮* _*PA*_ (the point pattern of presence/absence locations) are different, it doesn’t make sense that *𝒮* _*PO*_ and *𝒮* _*PA*_ follow the same model. In order to capture the influence of *𝒮* _*PO*_ on the *p*(**s**) surface associated with *𝒴* _*PA*_ (the presence/absence data, we could add *δ*_*PO*_*η*_*PO*_(**s**) to the mean for *Z*(**s**) in model (c) of Section 5.2, i.e. we could have a *δ*_*PA*_*η*_*PA*_(**s**) term and a *δ*_*PO*_*η*_*PO*_(**s**) term in order to improve prediction of presence/absence.

So, we have two sources for possible preferential sampling, one for each dataset. We might insist that *δ*_*PO*_ *>* 0. Then, from the presence-only data, the probability of presence will be increased around the 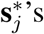 and decreased away from them. Indeed, the locations in *𝒮* _*PO*_ are severely biased; they are locations where we see only 1’s. We are severely over-sampling presences with *𝒮* _*PO*_ and we should increase probability of presence where we do.

In summary, we now have four potential models for *λ*_*PO*_(**s**), parallel to those for *λ*_*PA*_(**s**) to combine with the model for *Y* (**s**). Many of these models will be difficult to identify. We might focus our effort on a model for *𝒮* _*PO*_ analogous to model (ii) in Section 5.3 for *𝒮* _*PA*_. Then, we can add a *δ*_*PO*_*η*_*PO*_(**s**) term to the mean of *Z*(**s**) under (b), (c), or (d). In other words, the full model takes the form

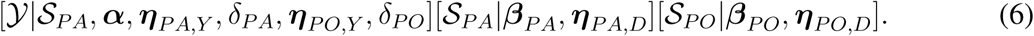

The Supplementary Material shows how to fit this model.

As a last remark, in practice, with a partial realization of the presence-only point pattern, we need to degrade *λ*_*PO*_(**s**) in the model fitting. The following subsection briefly reviews an approach to implement such degradation. The Supplementary Material shows how to adjust and fit (6) in the presence of a partially observed presence-only point pattern.

### 6.1 Spatial modeling of presence-only data in practice

Analysis of presence-only data has seen growth in recent years due to increased availability of such records from museum databases and other non-systematic surveys, see Graham et al. (2004). Presence-only data is not *inferior* to presence/absence data. In fact, it can be viewed as the opposite; in principle, presence-only data offer a complete census while presence/absence data, since confined to a specified set of sampling sites, contains less information. However, in practice, a complete census of individuals is rarely achieved. The sampling effort required to obtain such censuses usually exceeds the available resources.

An early model-based strategy for presence-only data attempts to implement a presence/absence approach by drawing so-called *background samples*, a random sample of locations in the region with known environmental features. These samples were characterized as pseudo-absences (Engler et al., 2004; Ferrier et al., 2002) and a logistic regression was fitted to the observed presences and these pseudo-absences, following Section 3. Since presence/absence is unknown for these samples, work of Pearce and Boyce (2006) and Ward et al. (2009) showed how to adjust the resulting logistic regression to account for this. In any event, this approach *manufactures* an arbitrary amount of data. Additionally, it ignores spatial dependence for presence/absence across locations. The observed presences, as a random number of random locations, should be viewed as a spatial point pattern (see Warton and Shepherd, 2010; Chakraborty et al., 2011, in this regard).

An algorithmic strategy in common use these days is the maximum entropy (Maxent) approach, (see, e.g., Phillips et al., 2006, 2009). Maxent is a constrained optimization method which finds the optimal species density (closest to a uniform) subject to moment constraints. The availability of an attractive software package (http://biodiversityinformatics.amnh.org/opensource/maxent/), encourages its use for presence-only data analysis. The resultant density surface is interpreted as providing the relative chance of observing a species at a given location compared to other locations in the region (and can not be interpreted as providing presence/absence probabilities). However, as an optimization strategy rather than a stochastic modeling approach, Maxent is unable to attach any uncertainty to resulting optimized estimates. Also, Maxent is unable to provide an intensity surface. Hence, for example, we are unable to determine the expected number of individuals in a specified region.

Arguably, a formal point pattern modeling approach is preferable since it enables full inference, with associated uncertainty, over the region. Modeling presence-only data as a point pattern specifies an associated intensity in terms of the available environments, at available spatial scale, across the region. Spatial structure for the intensity surface is introduced through spatial random effects, resulting in a log Gaussian Cox process (Møller et al., 1998; Møller and Waagepetersen, 2004), as discussed in Section 5 above.

Employing the LGCP in practice acknowledges that the observed point pattern is biased through anthropogenic processes, e.g., human intervention to transform the landscape and non-uniform (in fact, often very irregular) sampling effort. Such bias in sampling is a common problem, see for example Loiselle et al. (2008) and references therein. This requires adjusting the *potential* species intensity to a *realized* intensity which is treated as a *degradation* of the former. Such modeling adjustment is discussed in detail in Chakraborty et al. (2011) which we briefly review below.

Variation in site access is one of the factors that influences the likelihood of the site to be visited/sampled. For example, sites (i) adjacent to roads or along paths, (ii) near urban areas, (iii) with public ownership, e.g., state or national parks, or (iv) with flat topography are likely to be over-sampled relative to more inaccessible sites. When bias implies that only a portion of the region is sampled, it is likely that only a portion of the overall point pattern is observed. Land use, as a result of human intervention, affects *availability* of locations, hence, inference about the intensity. Also, agricultural transformation and dense stands of alien invasive species preclude availability. Transformed areas are not sampled and this information must also be included in the modeling. Altogether, sampling tends to be sparse and irregular; we rarely collect a random sample of available environments.

Detection can affect inference regarding the intensity. That is, we may incorrectly identify a species as present when it is actually absent (false presence) or fail to detect a species that is actually present (false absence) (Reese et al., 2005). Evidently, the prevalence of these false records will affect the performance of an explanatory model on response to environmental features (Tyre et al., 2003). Modeling for these errors can be attempted but requires information beyond the scope here.

### 6.2 Some explicit modeling details

Following ideas in Chakraborty et al. (2011), we conceptualize a *potential* intensity, i.e., the intensity in the absence of degradation, as well as a *realized* (or effective) intensity that operates in the presence of degradation. Further, we imagine that the intensity is tiled to grid cells at the resolution of the available environmental covariate surface. We imagine three surfaces over a region of interest, *D*. First, let *λ*(**s**) be the potential intensity surface, i.e., a positive function which is integrable over *D*. *λ*(**s**) is the intensity in the absence of degradation. With ∫ _*D*_ *λ*(**s**)*d***s** =*λ*(*D*), *g*(**s**) =*λ*(*s*)*/λ*(*D*) gives the potential density over *D*. Modeling for *λ*(**s**) is a log Gaussian Cox process (LGCP) which expects the environmental covariates, say **x**(**s**) to influence the intensity as a linear form in parameters. So, for any location *s ∈ D*, as in Section 5 above, we have

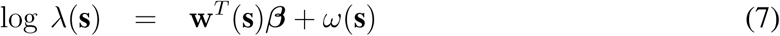

with *ω*(**s**), a zero-mean stationary, isotropic Gaussian process (GP) over *D*, to capture residual spatial association in the *λ*(**s**) surface across grid cells. With regard to preferential sampling, we propose to employ (33) as the model for an available presence-only dataset.

Turning to degradation, we envision an availability surface, *U* (**s**), a binary surface over *D* such that *U* (**s**) =1 or 0 according to whether location **s** is untransformed (hence, available) by land use or not. That is, assuming no sampling bias, *λ*(**s**)*U* (**s**) can only be *λ*(**s**) or 0 according whether **s** is available or not. Thirdly, we envision a sampling effort surface over *D* which we denote as *T* (**s**). *T* (**s**) is also a binary surface and *T* (**s**)*U* (**s**) =1 indicates that location **s** is both available and sampled. Altogether, *λ*(**s**)*U* (**s**)*T* (**s**) becomes the degraded intensity at location **s**. This implies that in regions where no locations were sampled, the operating intensity for the species is 0.

Suppose we partition *D* into grid cells with *A*_*i*_, *i* =1, 2, *…I* denoting the geographical region corresponding to grid cell *i*. Again, typically the gridding is at the resolution of the predictors used in explaining *λ*(**s**). Then, if we average *U* (**s**) over *A*_*i*_, we obtain 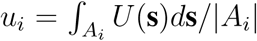 where *|A*_*i*_*|* is the area of cell *i*. Evidently, *u*_*i*_ is the proportion of cell *i* that is transformed. *u*_*i*_ can often be obtained, through remote sensing, for all grid cells. Further, we can set 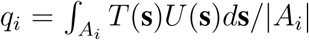 and interpret *q*_*i*_ as the probability that a randomly selected location in *A*_*i*_ was available and sampled. Thus, we can capture availability and sampling effort at areal unit scale. Additionally, 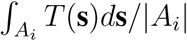 can be viewed as the sampling probability associated with cell *i*. Then, if *T* (**s**) is viewed as random, the expectation of the integral would yield 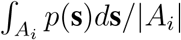 where, now, *p*(**s**) =*P* (*T* (**s**) =1) *∈* [0, 1]. Clearly, *p*(**s**) gives the local probabilities of sampling, not a probability density over *D*. Finally, if we define *p*_*i*_ through *q* _*i*_= *u* _*i*_ *p* _*i*_, then 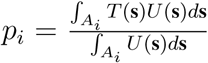, i.e., *p*_*i*_ is the conditional probability that a randomly selected location in cell *i* is sampled given it is available. As an illustration, we might set *p*_*i*_ equal to 1 or 0 which we interpret as *T* (**s**) =*U* (**s**) *∀***s** *∈ A*_*i*_ or *T* (**s**) =0 *∀***s** *∈ A*_*i*_, respectively. That is, either all available sites in *A*_*i*_ were visited or no available sites in *A*_*i*_ were visited. This degraded point pattern model is what we use for the data fusion. Fitting is described briefly in the Supplementary Material. Full details are provided in Chakraborty et al. (2011)

### 6.3 Model fitting and inference for data fusion

Here, we consider the following models for data fusion:

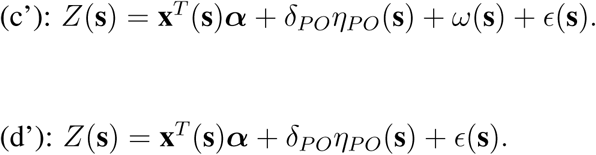

These models replace *δ*_*PA*_*η*_*PA*_(**s**) with *δ*_*PO*_*η*_*PO*_(**s**) in models (c) and (d). In addition to models (c’) and (d’), we consider two models which also include preferential sampling associated with the presence/absence data, *δ*_*PA*_*η*_*PA*_(**s**):

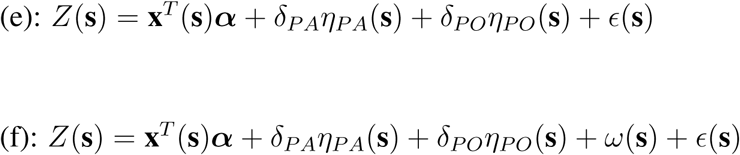

In the data fusion modeling, we assume an LGCP (model (ii) in Section 5.3) for *𝒮* _*PO*_, the set of presence locations of presence-only data. The same Bayesian priors and fitting as in Section 5.3 are applied for models (a)-(f) with details given in the Supplementary Material. As discussed in Section 6.2, we adopt the sampling effort surface *T* (**s**) for each grid cell so that *T* (**s**) =1 for all cells where at least one presence-only point is observed, *T* (**s**) =0 otherwise. The estimation and predictive performance results for models (a) and (b) are the same as those in Section 5.3. Since *δ*_*PO*_ is expected to be positive, a priori, we assume uniform prior on [0, 100], i.e., *δ*_*PO*_ ∼*𝒰* (0, 100).

Table 6 shows the estimation results for model (e) which include both *δ*_*PAηPA*_(**s**)+ *δ*_*POηPO*_(**s**). None of the *δ*_*PA*_ are significantly different from 0. However, all of the *δ*_*PO*_ are far from 0 revealing that the locations of the presence-only sites significantly improves the performance of the presenceabsence model. Table 7 shows the results for the TR measure under the same settings as in Section 5.3. The results are similar to those in Table 5. Performance is essentially indistinguishable across all models other than model (a) though model (f) emerges as the best. As a last remark here, if we focus on presence/absence locations which are near observed presence-only locations, we find an improvement in the TR measure for presence-absence at those locations compared to the corresponding model ignoring the presence-only data (results not shown).

**Table 6:** Estimation results for data fusion model (e).

| Model(e) | MR |  | OB |  | JB |  | GB |  |
| --- | --- | --- | --- | --- | --- | --- | --- | --- |
|  | Mean | 95% Int | Mean | 95% Int | Mean | 95% Int | Mean | 95% Int |
| const | -2.282 | [-2.670, -1.894] | -2.148 | [-2.560, -1.724] | -1.867 | [-2.298, -1.479] | -3.112 | [-4.153, -2.226] |
| mDR | 0.134 | [-0.160, 0.419] | 0.264 | [-0.088, 0.607] | 0.313 | [-0.036, 0.644] | 0.502 | [-0.196, 1.252] |
| maxTWM | 0.147 | [-0.185, 0.490] | -0.012 | [-0.419, 0.410] | -0.030 | [-0.431, 0.398] | -0.378 | [-1.310, 0.635] |
| meanTDQ | 0.745 | [0.280, 1.195] | 0.788 | [0.235, 1.343] | 0.659 | [0.112, 1.209] | 0.174 | [-0.930, 1.296] |
| minTCM | -0.023 | [-0.165, 0.123] | 0.052 | [-0.122, 0.230] | -0.066 | [-0.234, 0.094] | 0.525 | [0.107, 0.948] |
| PWM | 0.098 | [-0.046, 0.247] | 0.067 | [-0.108, 0.248] | 0.060 | [-0.116, 0.236] | -0.399 | [-0.881, 0.037] |
| PS | -0.207 | [-0.370, -0.042] | -0.077 | [-0.281, 0.123] | -0.128 | [-0.316, -0.060] | 0.033 | [-0.385, 0.442] |
| PWQ | -0.006 | [-0.179, 0.170] | -0.005 | [-0.228, 0.212] | 0.252 | [0.053, 0.453] | -0.041 | [-0.523, 0.438] |
| $\delta_{PA}$ | 0.034 | [-0.025, 0.093] | -0.030 | [-0.093, 0.027] | -0.003 | [-0.072, 0.063] | 0.107 | [-0.000, 0.231] |
| $\delta_{PO}$ | 0.463 | [0.362, 0.582] | 0.957 | [0.652, 1.449] | 0.831 | [0.654, 1.029] | 1.350 | [1.042, 1.905] |
| Model(e) | AO |  | BB |  | GM |  |  |  |
|  | Mean | 95% Int | Mean | 95% Int | Mean | 95% Int |  |  |
| const | -2.904 | [-3.632, -2.310] | -3.322 | [-4.061, -2.734] | -3.774 | [-4.977, -2.887] |  |  |
| mDR | 0.373 | [-0.052, 0.776] | -0.087 | [-0.474, 0.301] | 0.348 | [-0.197, 0.927] |  |  |
| maxTWM | -0.381 | [-0.866, 0.146] | 0.740 | [0.245, 1.259] | 0.061 | [-0.608, 0.726] |  |  |
| meanTDQ | 0.715 | [0.050, 1.343] | 0.158 | [-0.451, 0.790] | 1.065 | [0.127, 2.037] |  |  |
| minTCM | 0.032 | [-0.173, 0.229] | 0.050 | [-0.139, 0.238] | -0.024 | [-0.297, 0.253] |  |  |
| PWM | -0.127 | [-0.339, 0.087] | 0.128 | [-0.092, 0.356] | -0.393 | [-0.737, -0.062] |  |  |
| PS | -0.320 | [-0.577, -0.057] | -0.343 | [-0.603, -0.090] | -0.063 | [-0.397, 0.271] |  |  |
| PWQ | -0.368 | [-0.663, -0.093] | 0.270 | [-0.015, 0.564] | 0.601 | [0.225, 1.050] |  |  |
| $\delta_{PA}$ | -0.039 | [-0.118, 0.040] | 0.035 | [-0.043, 0.124] | 0.019 | [-0.098, 0.161] | | |
| $\delta_{PO}$ | 0.703 | [0.508, 0.933] | 0.817 | [0.587, 1.121] | 1.053 | [0.841, 1.342] | | |

**Table 7:** Estimation results for the TR measure for the data fusion. The results for (a) and (b) are the same as in Table 5

|  | Model (a) |  | Model (b) |  | Model (c) |  |
| --- | --- | --- | --- | --- | --- | --- |
|  | Mean | 95% Int | Mean | 95% Int | Mean | 95% Int |
| MR | 0.104 | [0.094, 0.114] | 0.168 | [0.145, 0.191] | 0.171 | [0.154, 0.193] |
| OB | 0.099 | [0.088, 0.109] | **0.183** | [0.163, 0.200] | 0.178 | [0.155, 0.198] |
| JB | 0.072 | [0.063, 0.081] | **0.201** | [0.180, 0.230] | 0.198 | [0.175, 0.223] |
| GB | 0.126 | [0.112, 0.139] | 0.405 | [0.372, 0.434] | 0.407 | [0.372, 0.436] |
| AO | 0.034 | [0.027, 0.043] | 0.095 | [0.068, 0.129] | 0.103 | [0.075, 0.139] |
| BB | 0.057 | [0.048, 0.068] | 0.112 | [0.078, 0.152] | 0.127 | [0.099, 0.163] |
| GM | 0.026 | [0.020, 0.032] | 0.135 | [0.111, 0.166] | 0.143 | [0.112, 0.182] |
|  | Model (d) |  | Model (e) |  | Model (f) |  |
|  | Mean | 95% Int | Mean | 95% Int | Mean | 95% Int |
| MR | 0.170 | [0.147, 0.195] | 0.165 | [0.136, 0.195] | **0.171** | [0.152, 0.197] |
| OB | 0.172 | [0.151, 0.197] | 0.169 | [0.134, 0.190] | 0.179 | [0.152, 0.207] |
| JB | 0.188 | [0.165, 0.216] | 0.186 | [0.159, 0.213] | **0.201** | [0.172, 0.221] |
| GB | 0.401 | [0.366, 0.440] | 0.404 | [0.363, 0.441] | **0.412** | [0.372, 0.449] |
| AO | 0.097 | [0.067, 0.137] | 0.101 | [0.073, 0.133] | **0.106** | [0.071, 0.144] |
| BB | 0.124 | [0.097, 0.158] | 0.124 | [0.085, 0.169] | **0.134** | [0.093, 0.169] |
| GM | 0.144 | [0.101, 0.189] | 0.150 | [0.102, 0.195] | **0.156** | [0.115, 0.197] |

## 7 Summary and future work

Our contribution is to attempt to bring more clarity to a frequent activity for ecologists, modeling presence/absence. We have done this from a probabilistic perspective, arguing that presence/absence data should be viewed as a point level phenomenon and therefore, stochastic modeling for presence/absence should be done at unitless points. In the development we have also argued that attempting to model presence/absence at areal scale raises challenges and, further, that any such modeling is incompatible with point-level modeling. Continuing, we have asserted that the number of presences in a fixed bounded region must be finite and therefore, that a physical realization of a presence in the region is larger than a point.

We acknowledge that we are being more formal in developing this perspective than is customary. When presence/absence data is supplied at point-level, it will be a geo-coded location and, often it is supplied at areal scale. However, this is not a deterrent from using our perspective. All continuous measurements are obtained up to rounding error. When a temperature is recorded at a location, the location is provided up to the accuracy of the geo-coding device; nonetheless, we routinely model temperatures at (unitless) points.

Next, we turned our attention to improving modeling for a presence/absence dataset. We introduced the utility of preferential sampling in this context, anticipating that there may be bias in sampling sites visited for presence/absence data; sampling may favor seeing more presences. We argued that the idea of a shared process model, viewing the set of presence/absence locations as a point pattern, can improve inference regarding the presence/absence surface. We demonstrated this with a plant presence/absence dataset from New England.

Finally, we asserted that presence-only data should be modeled as a point pattern, albeit degraded due to availability and sampling effort. We argued that this makes it evident that a common model for presence/absence and for presence-only data can not be stochastically coherent. Hence, if we seek a data fusion having both presence/absence data as well as presence-only data, a different approach is needed. We argued that, again, a shared process specification is coherent for such fusion and illustrated by adding a presence-only plant dataset from New England to the previous presence/absence dataset.

Future work offers much opportunity. More experience is needed with regard to the rich set of modeling specifications that we have presented in Sections 5 and 6. We also anticipate the need to supply user-friendly software to enable ecologists to play with these models with their own datasets. A particularly useful future direction leads us to joint species distribution models. These are easy to envision but challenging to fit. Another useful future direction will consider different types of response data, e.g., abundance or basal area, where again, preferential sampling of locations may occur.

## Acknowledgement

The authors thank John Silander for numerous useful conversations which motivated much of the formalism presented here. They also thank Jenica Allen for supplying the IPANE and GBIF datasets which illustrated our ideas. The computational results were obtained using Ox version 6.21 (Doornik, 2007).

## A. Further exploratory analysis of the data

Figure 4 shows the standardized covariate surfaces for the 7 selected covariates. The location which indicates extreme values in maxTWM and PWM corresponds to the summit of Mt. Washington which is notorious for exhibiting extreme climate conditions.

**Figure 4:**
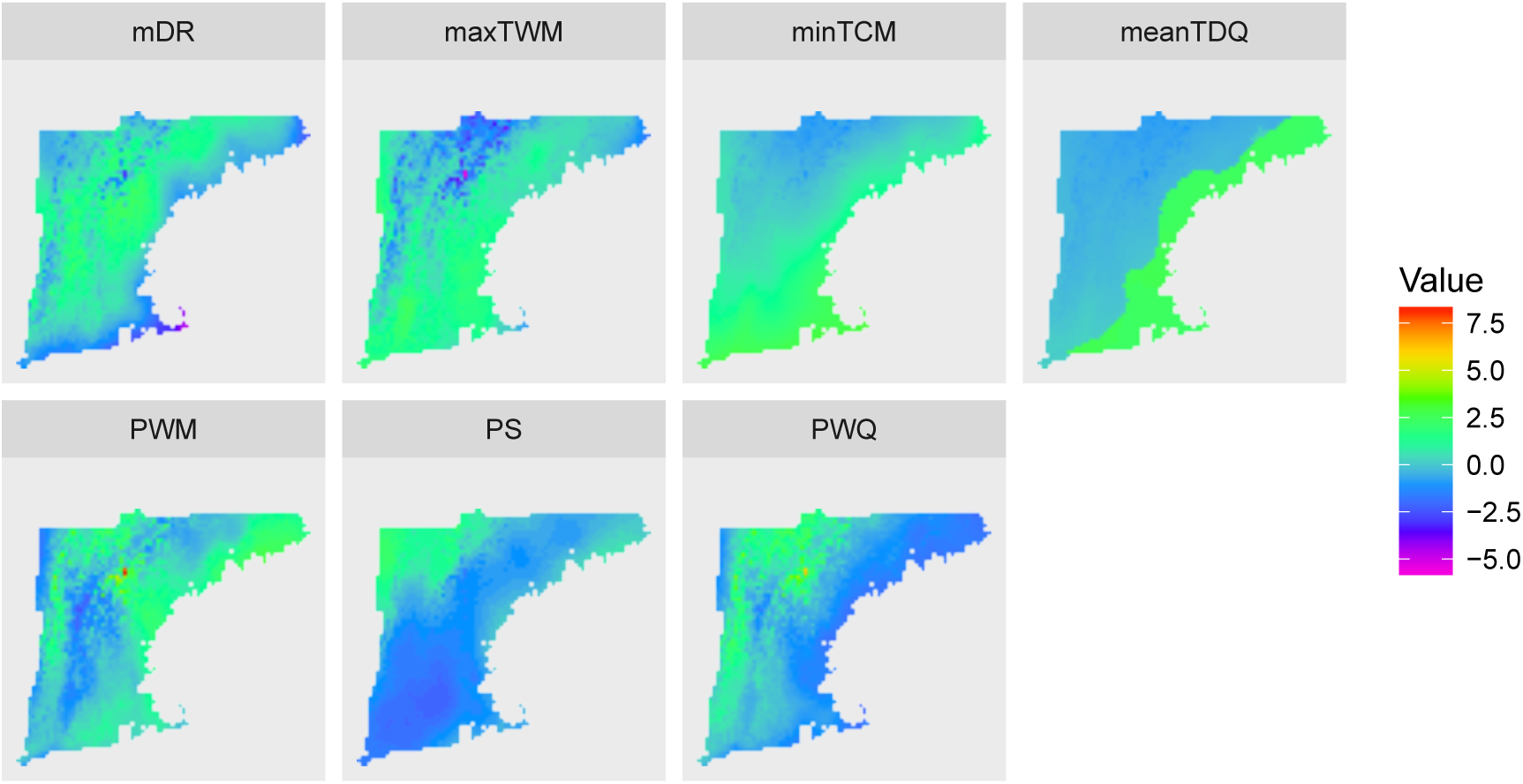
The standardized covariate surface for selected 7 covariates across the study region.

We also show the estimation results of the LGCP for *𝒮* _*PA*_. Bayesian inference of this model is described in Section B.1. We adopt weak prior specification: 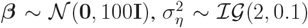 and *ϕ*_*η*_ ∼*𝒰* (0, 200). We discard the first 20,000 samples as burn-in and preserve the subsequent 20,000 samples as posterior samples. Table 8 shows the estimation results for the parameters in the LGCP model. All covariates are significant except for mDR. Figure 5 shows the posterior mean surface for ***η*** and log *λ*.

**Figure 5:**
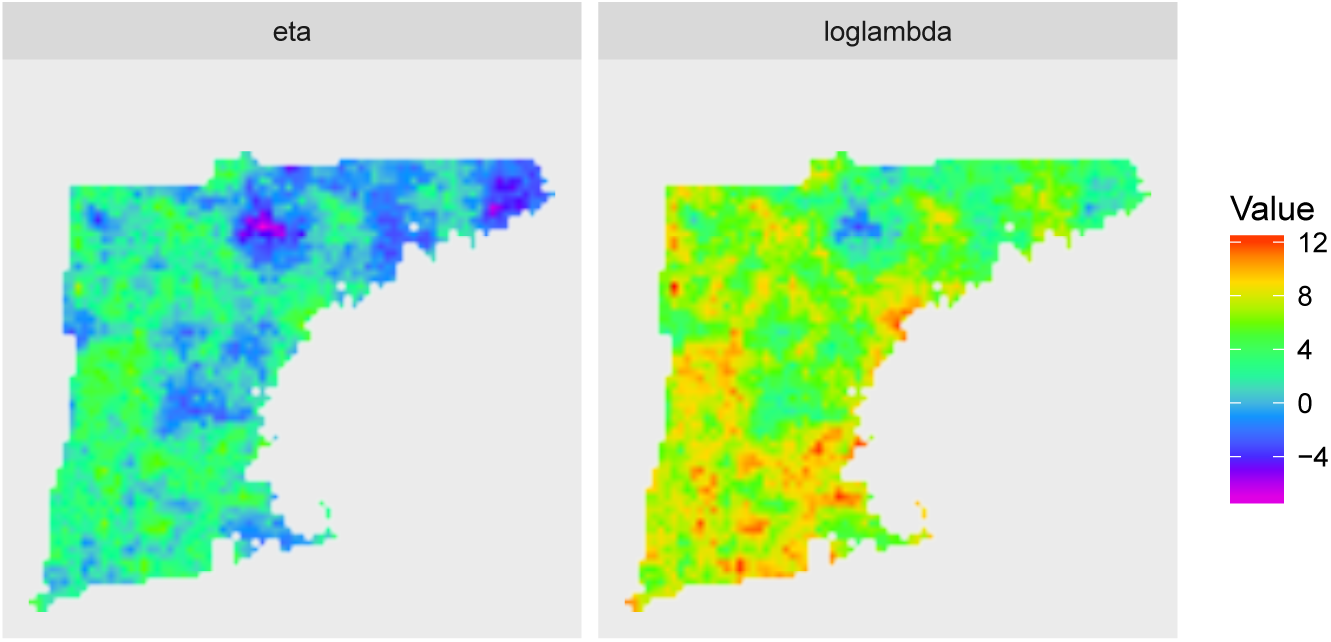
The posterior mean surface for ***η*** (left) and log ***λ*** (right) for *𝒮* _*PA*_.

## B. Model fitting details

### B.1. Model fitting details for presence/absence data with preferentail sampling

Models for *Z*(**s**) are discussed in Section 5. Let *µ*(**s**) be the main effect for *Z*(**s**), i.e., *Z*(**s**) =*µ*(**s**) + *c*(**s**) where *c*(**s**) ∼ *N* (0, *τ* ^2^). The specification for *µ*(**s**) depends on the model, e.g., *µ*(**s**) =**x**^*T*^ (**s**)***α*** + *δη*(**s**) for the preferential sampling model (d) in Table 2 of our manuscript. We denote ***Z*** =(*Z*(**s**_1_), *Z*(**s**_2_), *…, Z*(**s**_*n*_))^*T*^, ***µ*** =(*µ*(**s**_1_), *µ*(**s**_2_), *…, µ*(**s**_*n*_))^*T*^ and **X** =(**x**(**s**_1_), **x**(**s**_2_), *…*, **x**(**s**_*n*_))^*T*^.

As for prior specification, we assume 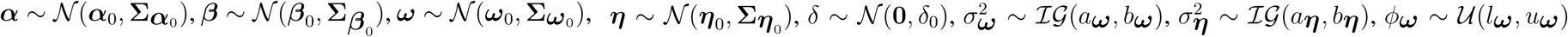 and *ϕ***_*η*_** ∼*𝒰* (*l***_*η*_**, *u***_*η*_**). The likelihood for *𝒮* _*PA*_ is

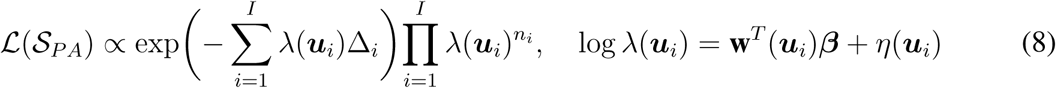

Where 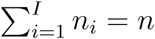.

**Gibbs sampling for *Z***(*·*)**: Model (a)-(d)**

- Sampling *Z*(**s**_*i*_) ∼*𝒯 𝒩* _*>*0_(*µ*(**s**_*i*_), *τ* ^2^) when *Y* (**s**_*i*_) =1 and *Z*(**s**_*i*_) ∼*𝒯 𝒩* _*≤*0_(*µ*(**s**_*i*_), *τ* ^2^) otherwise for *i* =1, 2, *…, n* where *𝒯 𝒩* _*>*0_ (*𝒯 𝒩* _*≤*0_) denotes a truncated normal distribution on positive (nonpositive) domain.

**Gibbs sampling for *α*: Model (a)-(d)**

- Sampling ***α*** ∼*𝒩* (***µ*_*α*_**, **Σ_*α*_**)

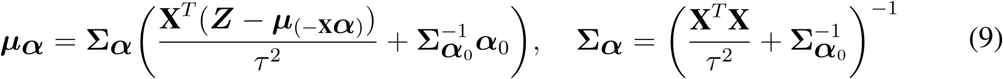

where ***µ***_(-**X*α***)_ is ***µ*** except for **X*α***.

**Metropolis-Hastings update for *β*: Model (c) and (d)**

- MH algorithm for ***β***

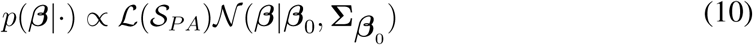

**Gibbs sampling for *ω*: Model (b) and (c)**

- Sampling ***ω*** ∼*𝒩* (***µ*_*ω*_**, **Σ_*ω*_**)

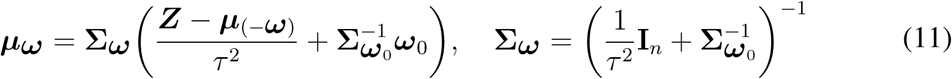

where ***µ***_(-***ω***)_ is ***µ*** except for ***ω***.

**Metropolis-Hastings update for *η*: Model (c) and (d)**

- MH algorithm for ***η***

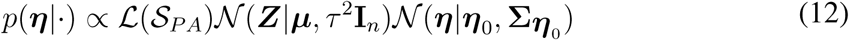

**Metropolis-Hastings update for 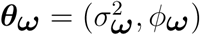: Model (b) and (c)**

- **MH algorithm for *θ*_*ω*_**

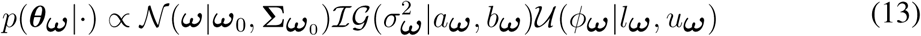

**Metropolis-Hastings update for 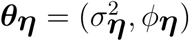: Model (c) and (d)**

- **MH algorithm for** *θ*_*η*_

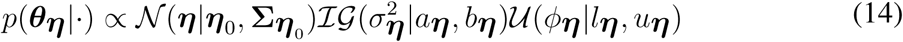

**Gibbs sampling for** *δ***: Model (c) and (d)**

- Sampling *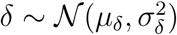*

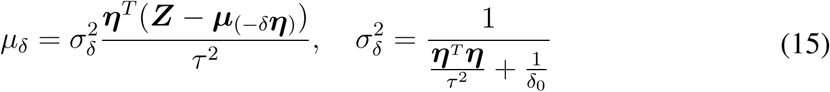

where ***µ***_(*-δ****η***)_ is ***µ*** except for *δ****η***.

For large *n*, we implement nearest neighbor Gaussian processes (NNGP, Datta et al., 2016). We explain for ***ω*** below, but the same discussion can be applied for ***η***. NNGP expresses the joint density of ***ω*** as the product of approximate conditional densities by projecting on *neighbors* instead of the full set of locations, i.e.,

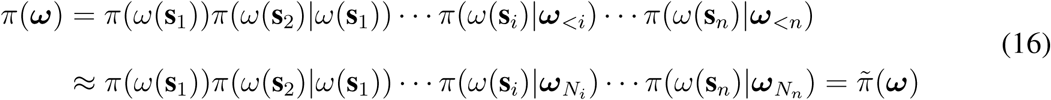

where ***ω***_*<i*_ ={*ω*(**s**_1_), *ω*(**s**_2_), *…, ω*(**s**_*i-*1_)} and *N*_*i*_ is the set of indices of neighbors of 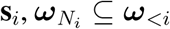 (see, e.g., Vecchia (1988), Stein et al. (2004), Gramacy et al. (2014) and Gramacy and Apley (2015)). 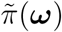 is a proper multivariate joint density (Datta et al. (2016)). As for neighbor selections, choosing *N*_*i*_ to be any subset of {1, 2, *…, i −* 1} ensures a valid probability density. For example, Vecchia (1988) specified *N*_*i*_ to be the *k* nearest neighbors of **s**_*i*_ among {1, 2, *…, i −* 1} with respect to Euclidean distance. Sampling from 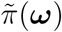 is sequentially implemented for *i* =1, *…, n* as follows

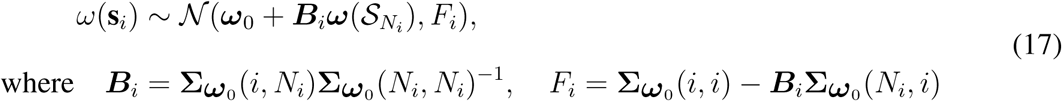

where ***B***_*i*_ =**Σ_*ω*0_**(*i, N*_*i*_)**Σ _*ω*0_** (*N*_*i*_, *N*_*i*_)^−1^, *F*_*i*_ =**Σ _*ω*0_** (*i, i*) − ***B***_*i*_**Σ _*ω*0_** (*N*_*i*_, *i*)

where 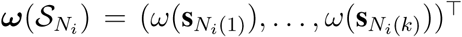 and *N*_*i*_(*j*) is *j*-th component of *N*_*i*_. Given this expression, Gibbs sampling for ***ω*** is available within the generalized spatial linear model framework (Datta et al. (2016)).

From the matrix perspective, we can write the multivariate Gaussian density *𝒩* (***ω****|****ω***_0_, **Σ**_*ω*0_) as a linear model

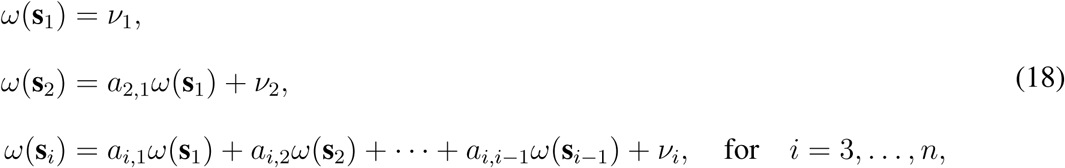

simply as ***ω*** =**A*ω*** + ***𝓋*** where **A** is *n× n* strictly lower-triangular with elements *a*_*i,j*_ =0 whenever *j ≥ i* and ***𝓋*** ∼***𝒩*** (***ω***_0_, **V**) where **v** is a diagonal matrix with *v*_*i,i*_ =*Var*[*ω*(**s**_*i*_)*|****ω***_*<i*_]. It is obvious that **I** − **A** is nonsingular and **Σ_*ω*0_**= (**I** − **A**)^−1^**V**(**I** − **A**)^*−T*^. The neighbor selection corresponds to introducing sparsity into **A**, i.e., *a*_*i,j*_ ≠ 0 when 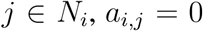 otherwise. The approximated covariance matrix is obtained as 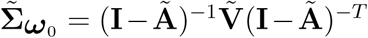 where 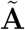 is a sparse approximation of **A** and the diagonal component of 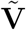 is 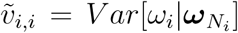. This can be performed in order *𝒪* (*nk*^3^) (linear in *n*) and in parallel across rows of **A** (Datta et al., 2016).

### Model fitting details for the presence/absence presence-only data fusion

Essentially the same discussion as in Section B.1 can be applied for the data fusion models (c’)-(f) in Section 6.3 in our manuscript. As for prior specification, we assume 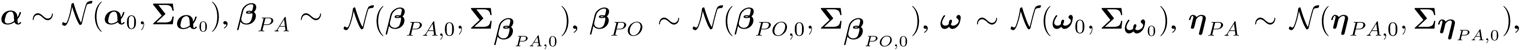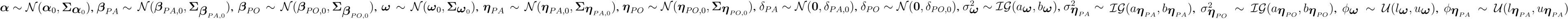, and 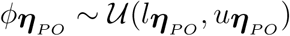. The likelihoods for *𝒮*_*PA*_ and *𝒮*_*PO*_ are

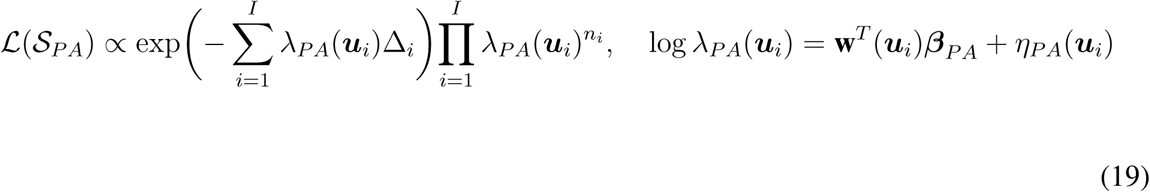

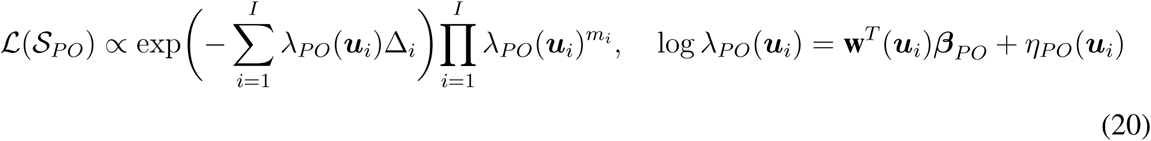

Where 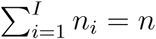 and 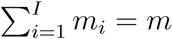.

**Gibbs sampling for *Z***(*·*)**: Model (c’)-(f)**

- Sampling *Z*(**s**_*i*_) ∼*𝒯 𝒩* _*>*0_(*µ*(**s**_*i*_), *τ* ^2^) when *Y* (**s**_*i*_) =1 and *Z*(**s**_*i*_) ∼*𝒯 𝒩*_*≤*0_(*µ*(**s**_*i*_), *τ* ^2^) otherwise for *i* =1, 2, *…, n* where *𝒯 𝒩* _*>*0_ (*𝒯 𝒩*_*≤*0_) is truncated normal distribution on positive (nonpositive) domain.

**Gibbs sampling for *α*: Model (c’)-(f)**

- Sampling ***α*** ∼*𝒩* (***µ*_*α*_**, **Σ_*α*_**)

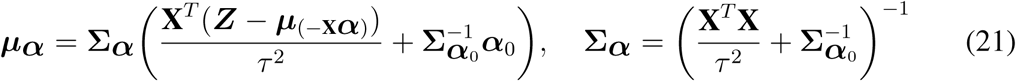

where ***µ***_(-**X*α***)_ is ***µ*** except for **X_*α*_**.

**Metropolis-Hastings update for *β***_*PA*_**: Model (e) and (f)**

- MH algorithm for ***β***_*PA*_

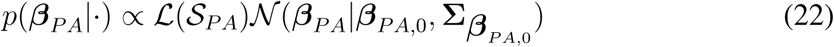

**Metropolis-Hastings update for *β***_*PO*_ **: Model (c’)-(f)**

- MH algorithm for ***β***_*PO*_

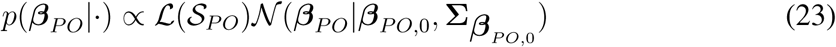

**Gibbs sampling for *ω*: Model (c’) and (f)**

- Sampling ***ω*** ∼*𝒩* (***µ*_*ω*_**, **Σ_*ω*_**)

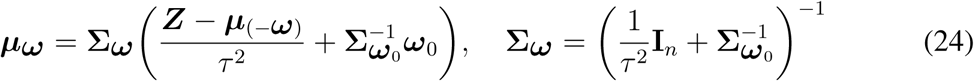

where ***µ***_(-***ω***)_ is ***µ*** except for ***ω***.

**Metropolis-Hastings update for *η***_*PA*_**: Model (e) and (f)**

- MH algorithm for ***η***_*PA*_

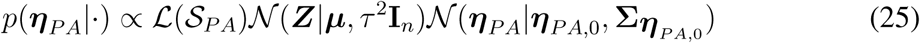

**Metropolis-Hastings update for *η***_*PO*_ **: Model (c’)-(f)**

- MH algorithm for ***η***_*PO*_

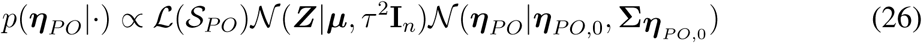

**Metropolis-Hastings update for 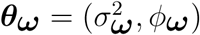: Model (b) and (c’)**

- MH algorithm for *θ*_*ω*_

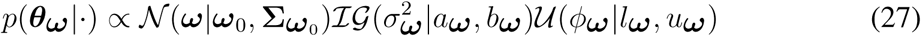

**Metropolis-Hastings update for** 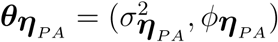: **Model (e) and (f)**

- MH algorithm for 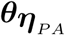

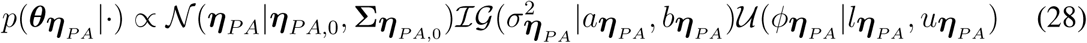

**Metropolis-Hastings update for** 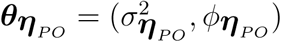: **Model (c’)-(f)**

- MH algorithm for 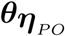

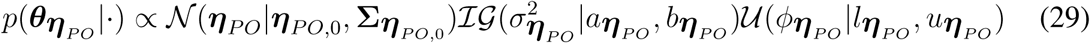

**Gibbs sampling for *δ*_*PA*_: Model (e) and (f)**

- Sampling 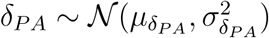

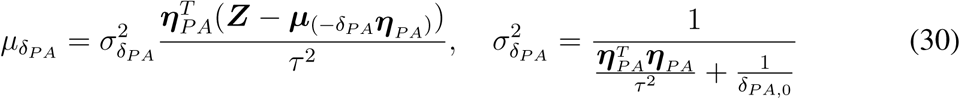

where 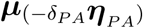 is ***µ*** except for *δ*_*PA*_***η***_*PA*_.

**Gibbs sampling for *δ*_*PO*_: Model (c’)-(f)**

- Sampling 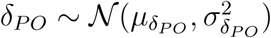

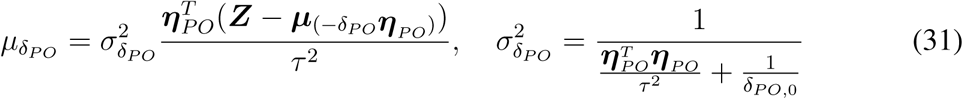

where 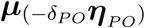 is ***µ*** except for *δ*_*PO*_***η***_*PO*_.

As discussed in Section B.1, we can implement NNGP completion, 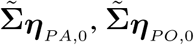 and 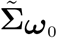.

### B.3. Model fitting for partially observed presence-only data

Under the gridding in Section 5, suppose we have *n*_*i*_ presence locations 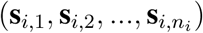 within grid cell *i* for *i* =1, 2, *…, I*. Following the discussion above, *U* (**s**_*i,j*_)*T* (**s**_*i,j*_) ≡ 1, 0 *≤ j ≤ n*_*i*_, 1 *≤ i ≤ I*. Then the likelihood function corresponding becomes

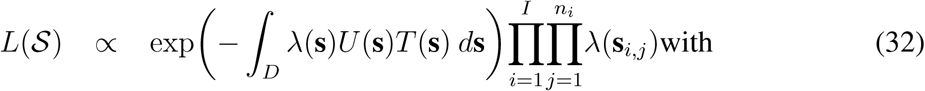

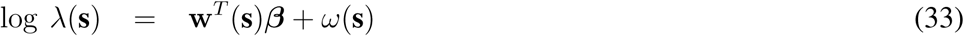

and *ω*(**s**), a zero-mean stationary, isotropic Gaussian process (GP) over *D*, to capture residual spatial association in the *λ*(**s**) surface across grid cells. Although we have only finitely many presence locations, the integral in *L*(*S*) involves the uncountable random field *λ*_*D*_ ={*λ*(**s**): **s** ∈ *D*}. Fortunately, we have a natural approximation to the stochastic integral at the scale of grid cells. That is, though we have geo-coded locations for the observed sites, with covariate information at grid cell level, we only attempt to explain the point pattern at grid cell level. Also, with many unsampled cells, many *n*_*i*_ =0.

A computational advantage accrues to working at grid cell level; we can employ a product Poisson likelihood approximation rather than the point pattern likelihood in (32). That is, for cell *i*, suppose **s**_*i*_ is the centroid. Then, given the set {*λ*(**s**_*i*_), *i* =1, 2, *…, I*, the *n*_*i*_ are independent and *n*_*i*_ ∼ Po(△*λ*(**s**_*i*_)*q*_*i*_) where △ is the area of cell *i*. Approximation of the point pattern likelihood using such a *tiled* surface over a lattice embedding the region was discussed in Benes et al. (2002). There it is shown that the approximation can be justified in the sense that the resulting approximate posterior converges to the true posterior as the partition gets finer.

Notice that, for any cell with *q*_*i*_ = 0 (which can happen if either *p*_*i*_ = 0 or *u*_*i*_ = 0) there is no contribution from *A*_*i*_ in the product Poisson likelihood. Let **W** = (**w**(**s**_1_), **w**(**s**_2_), *…*, **w**(**s**_*m*_))^*T*^ Since, from (33), log*λ*(**s**) follows a GP, the posterior distribution takes the form

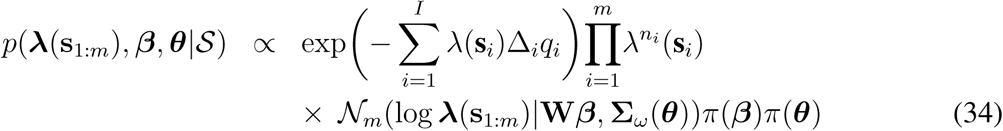

where *N*_*m*_(*·|·*) denotes an *m* (*< I*) dimensional Gaussian density with mean **W*β*** and covariance **Σ**_*ω*_(***θ***) and ***θ*** denotes the parameters in the covariance function of *ω*(**s**) in (33).

With regard to inference, posterior samples of ***β*** help us to infer whether a particular factor has a significant impact (positive or negative) on species intensity. The *ϕ* parameter indicates the strength of spatial association for the realization of the intensity surface over *D*. This association may arise because some potentially important covariates are not available or because the covariate impact is not well captured using a linear form. That is, since Gaussian processes can capture a wide range of dependencies, using them in a hierarchical setting enhances predictive performance for the model.

With regard to displays of intensity surfaces, the *λ*_*i*_*p*_*i*_ surface will capture the (lack of) sampling effort. The *λ*_*i*_*u*_*i*_ surface reveals the effect of transformation. Of course, the *λ*(**s**) surface is most interesting since it offers insight into the expected pattern of presences over all of *D*. Posterior draws of *λ*_1:*I*_ can be used to infer about the potential intensity, displaying say the posterior mean surface. We can also learn about the potential density *g*(**s**) in this discretized setting as 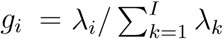, and the corresponding density under transformation as 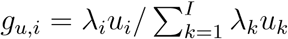.

**Table 8:** Estimation results of LGCP for *𝒮* _*PA*_

|  | Mean | Stdev | 95% Int |  | Mean | Stdev | 95% Int |
| --- | --- | --- | --- | --- | --- | --- | --- |
| const | 7.687 | 0.139 | [7.452, 7.984] | mDR | -0.019 | 0.159 | [-0.283, 0.306] |
| maxTWM | 0.538 | 0.171 | [0.101, 0.816] | minTCM | 1.913 | 0.131 | [1.618, 2.135] |
| meanTDQ | 0.263 | 0.119 | [0.020, 0.472] | PWM | -0.510 | 0.107 | [-0.713, -0.263] |
| PS | 0.589 | 0.134 | [0.375, 0.852] | PWQ | 1.287 | 0.091 | [1.115, 1.493] |
| $\sigma_\eta^2$ | 5.713 | 0.291 | [5.145, 6.289] | $\phi_\eta$ | 0.507 | 0.030 | [0.450, 0.553] |

## Footnotes

1 To expedite the flow we defer referencing to subsequent sections.

2 We accept this idea routinely in looking at data. We never observe continuous measurements; we only observe them up to decimal accuracy. Nonetheless, conceptually, we proceed to model them as continuous.

3 Even for an abundant species where the number of individuals may be so large that we might want to think of the number as countably infinite, formally we still have the same issues as above.

4 With second stage modeling, as discussed in Section 3, we also introduce a latent Gaussian process, *Z*(**s**).

## References

Aarts, G., J. Fieberg, and J. Matthiopoulos (2012). Comparative interpretation of count, presence-absence and point methods for species distribution models. Methods in Ecology and Evolution 3, 177–187.

Banerjee, S., B. P. Carlin, and A. E. Gelfand (2014). Hierarchical Modeling and Analysis for Spatial Data, 2nd ed. Chapman and Hall/CRC.

Benes, V., K. Bodlák, J. Møller, and R. P. Waagepetersen (2002). Bayesian analysis of log Gaussian Cox processes for disease mapping. Technical report, Department of Mathematical Sciences, Aalborg University.

Besag, J. E. (1974). Spatial interaction and the statistical analysis of lattice systems. Journal of the Royal Statistical Society, Series B 36, 192–236.

Cecconi, K., L. Grisotto, D. Catelan, C. Lagazio, V. Berrocal, and A. Biggeri (2016). Preferential sampling and Bayesian geostatistics: statistical modeling and examples. Statistical Methods in Medical Research 25, 1224–1243.

Chakraborty, A., A. E. Gelfand, A. M. Wilson, A. M. Latimer, and J. A. Silander (2011). Point pattern modelling for degraded presence-only data over large regions. Journal of the Royal Statistical Society, Series C 60, 757–776.

Clark, J. S., D. Nemergut, B. Seyednasrollah, P. Turner, and S. Zhange (2017). Generalized joint attribute modeling for biodiversity analysis: median-zero, multivariate, multifarious data. Ecological Monographs 87, 34–56.

Cressie, N. and C. K. Wikle (2011). Statistics for Spatio-Temporal Data. Wiley.

Datta, A., S. Banerjee, A. O. Finley, and A. E. Gelfand (2016). Hierarchical nearest-neighbor Gaussian process models for large geostatistical datasets. Journal of the American Statistical Association 111, 800–812.

Diggle, P., R. Menezes, and T. Su (2010). Geostatistical inference under preferential sampling. Journal of the Royal Statistical Society, Series C 59, 191–232.

Diggle, P. J., J. A. Tawn, and R. A. Moyeed (1998). Model-based geostatistics. Journal of the Royal Statistical Society, Series C 47, 299–350.

Doornik, J. (2007). Ox: Object Oriented Matrix Programming. Timberlake Consultants Press.

Dorazio, R. M. (2014). Accounting for imperfect detection and survey bias in statistical analysis of presence-only data. Global Ecology and Biogeography 23, 1472–1484.

Elith, J., C. Graham, R. Anderson, M. Dudik, S. Ferrier, A. Guisan, R. J. Hijmans, F. Huettmann, J. R. Leathwick, A. Lehmann, J. Li, L. G. Lohmann, B. A. Loiselle, G. Manion, C. Moritz, M. Nakamura, Y. Nakazawa, A. T. Peterson, S. J. Phillips, K. Richardson, R. Scachetti-Pereira, R. E. Schapire, J. Soberón, S. Williams, M. S. Wisz, and N. E. Zimmermann (2006). Novel methods improve prediction of species’ distributions from occurrence data. Ecography 29, 129–151.

Engler, R., A. Guisan, and L. Rechsteiner (2004). An improved approach for predicting the distribution of rare and endangered species from occurrence and pseudo-absence data. Journal of Applied Ecology 41, 263–274.

Ferrier, S., G. Watson, J. Pearce, and M. Drielsma (2002). Extended statistical approaches to modelling spatial pattern in biodiversity in northeast New South Wales. I. Species-level modelling. Biodiversity and Conservation 11, 2275–2307.

Fithian, W., J. Elith, T. Hastie, and D. A. Keith (2015). Bias correction in species distribution models: pooling survey and collection data for multiple species. Methods in Ecology and Evolution 6, 424–438.

Giraud, C., C. Calenge, and R. Julliard (2016). Capitalising on opportunistic data for monitoring biodiversity. Biometrics 72, 649–658.

Graham, C. H., S. Ferrier, F. Huettman, C. Moritz, and A. T. Peterson (2004). New developments in museum-based informatics and applications in biodiversity analysis. Trends in Ecology and Evolution 19, 497–503.

Gramacy, R. B. and D. W. Apley (2015). Local Gaussian process approximation for large computer experiments. Journal of Computational and Graphical Statistics 24, 561–578.

Gramacy, R. B., J. Niemi, and R. M. Weiss (2014). Massively parallel approximate Gaussian process regression. SIAM/ASA Journal of Uncertainty Quantification 2, 564–584.

Guisan, A., J. T C Edwards, and T. Hastie (2002). Generalized linear and generalized additive models in studies of species distributions: setting the scene. Ecological Modelling 157, 89–100.

Hastie, T. and W. Fithian (2013). Inference from presence-only data; the ongoing controversy. Ecography 36, 864–867.

Illian, J., A. Penttinen, H. Stoyan, and D. Stoyan (2008). Statistical Analysis and Modelling of Spatial Point Patterns. Wiley.

Loiselle, B. A., P. Jorgensen, T. Consiglio, I. Jimenez, J. G. Blake, L. G. Lohmann, and O. M. Montiel (2008). Evaluating plant collection representation for ecological niche modeling: a case study using plant vouchers from Ecuador and Bolivia. Journal of Computational and Graphical Statistics 35, 105–116.

Møller, J., A. R. Syversveen, and R. P. Waagepetersen (1998). Log Gaussian Cox processes. Scandinavian Journal of Statistics 25, 451–482.

Møller, J. and R. Waagepetersen (2004). Statistical Inference and Simulation for Spatial Point Processes. Chapman and Hall/RC.

Nychka, D. and J. Anderson (2010). Data assimilation. In Handbook of Spatial Statistics, pp. 477–492. CRC/Chapman and Hall.

Ovaskainen, O., D. B. Roy, R. Fox, and B. J. Anderson (2016). Uncovering hidden spatial structure in species communities with spatially explicit joint species distribution models. Methods in Ecology and Evolution 7, 428–436.

Pacifici, K., B. J. Reich, D. A. W. Miller, B. Gardner, G. Stauffer, S. Singh, A. McKerrow, and J. A. Collazo (2017). Integrating multiple data sources in species distribution modeling: a framework for data fusion. Ecology, 840–850.

Pati, D., B. J. Reich, and D. B. Dunson (2011). Bayesian geostatistical modelling with informative sampling locations. Biometrika 98, 35–48.

Pearce, J. L. and M. S. Boyce (2006). Modelling distribution and abundance with presence-only data. Journal of Applied Ecology 43, 405–412.

Phillips, S. J., R. P. Anderson, and R. E. Schapire (2006). Maximum entropy modeling of species geographic distributions. Ecological Modelling 190, 231–259.

Phillips, S. J., M. Dudík, J. Elith, C. H. Graham, A. Lehmann, J. Leathwick, and S. Ferrier (2009). Sample selection bias and presence-only distribution models: implications for background and pseudo-absence data. Ecological Applications 19, 181–197.

Reese, G. C., K. R. Wilson, J. A. Hoeting, and C. H. Flather (2005). Factors affecting species distribution predictions: a simulation modeling experiment. Ecological Applications 15, 554–564.

Royle, J. A., R. B. Chandler, C. Yackulic, and J. D. Nichols (2012). Likelihood analysis of species occurrence probability from presence-only data for modelling species distributions. Methods in Ecology and Evolution 3, 545–554.

Rundel, C. W., E. M. Schliep, A. E. Gelfand, and D. M. Holland (2015). A data fusion approach for spatial analysis of speciated *PM* 2.5 across time. Environmetrics 26, 515–525.

Sahu, S. K., A. E. Gelfand, and D. M. Holland (2016). Fusing point and areal level space-time data with application to wet deposition. Journal of the Royal Statistical Society, Series C 59, 77–103.

Saltzman, N. and D. Nychka (1998). DI, a design interface for constructing and analyzing spatial designs. Springer Science and Media.

Stein, M., Z. Chi, and L. Welty (2004). Approximating likelihoods for large spatial data sets. Journal of the Royal Statistical Society, Series B 66, 275–296.

Tjur, T. (2009). Coefficients of determination in logistic regression models-A new proposal: the coefficient of discrimination. The American Statistician 63, 366–372.

Tyre, A. J., B. Tenhumberg, S. A. Field, D. Niejalke, K. Parris, and H. P. Possingham (2003). Improving precision and reducing bias in biological surveys: estimating false-negative error rates. Ecological Applications 13, 1790–1801.

Vecchia, A. V. (1988). Estimation of model identification for continuous spatial processes. Journal of the Royal Statistical Society, Series B 50, 297–312.

Ver Hoef, J. M., N. Cressie, R. N. Fisher, and T. J. Case (2001). Uncertainty and spatial linear models for ecological data. Spatial Uncertainty in Ecology 69, 275–294.

Ward, G., T. Hastie, S. Barry, J. Elith, and J. Leathwick (2009). Presence-only data and the EM algorithm. Biometrics 65, 554–563.

Warton, D. I. and L. C. Shepherd (2010). Poisson point process models solve the pseudo-absence problem for presence-only data in ecology. Annals of Applied Statistics 4, 1383–1402.

Wikle, C. K. and L. M. Berliner (2007). A Bayesian tutorial for data assimilation. Physica D 230, 1–16.

Wikle, C. K., R. F. Milliff, D. Nychka, and L. M. Berliner (2001). Spatiotemporal hierarchical Bayesian modeling: tropical ocean surface winds. Journal of the American Statistical Association 96, 382–397.

